# Global analysis of membrane protein S-acylation in the model plant *Arabidopsis thaliana*

**DOI:** 10.1101/2025.07.01.658617

**Authors:** Lijuan Zhou, Lingtao Su, Mowei Zhou, Marina A. Gritsenko, Jinrong Wan, Yahui Ma, Yue Zhao, Ljiljana Paša-Tolić, Dong Xu, Gary Stacey

**Author notes:** Authors for correspondence: *Gary Stacey,* *Lijuan Zhou,. Department of Chemistry, Zhejiang University, Hangzhou, Zhejiang 310058, China (Mowei Zhou).

## Abstract

Protein S-acylation is the addition of fatty acids to the cysteine residues in a protein, catalyzed by protein S-acyltransferases (PATs). Despite extensive research on protein S-acylation in animals, our understanding of this process in plants remains limited. In this study, we sought to characterize the S-acylproteome of membrane proteins in *Arabidopsis* and identify potential substrates for two important plant immunity-related PATs (PAT5 and PAT9). To achieve this, S-acylated membrane proteins were first enriched via our optimized acyl-biotinyl exchange strategies at both the protein-level and peptide-level. The enriched samples were then analyzed by label-free quantitative liquid chromatography-mass spectrometry. The results from the two enrichment methods demonstrated that they were complementary in identifying S-acylated proteins and S-acylation sites. Using these methods, over 2500 S-acylation sites in more than 2000 putative S-acylated proteins were identified. Proteins involved in vesicle trafficking, plant phosphorylation, immune responses, and signal transduction pathways were significantly enriched. Additionally, certain amino acid patterns surrounding the S-acylation sites were identified. Comparisons of the S-acylproteomes between the wild type and the PAT5 and PAT9 mutants revealed over 100 potential substrates for both S-acyltransferases. The high quality of our data was supported by the significant overlap with the previously reported data and successful experimental verification of selected candidate proteins. Overall, our study revealed a well-represented S-acylproteome for *Arabidopsis* (especially its membrane proteins) and identified potential substrates for PAT5 and PAT9. These findings will facilitate the functional characterization of S-acylated proteins in plants.

## Introduction

Protein S-acylation is an essential post-translational modification (PTM) that involves the covalent attachment of a long fatty acid chain to specific cysteine residues in proteins through a thioester bond. Since the most commonly attached fatty acid is the 16-carbon saturated palmitic acid, S-acylation is often called S-palmitoylation or palmitoylation. Different from the other protein lipid modifications, protein S-acylation is reversible in cells, controlled by protein S-acyltransferases (PATs) and de-S-acylation enzymes (Dekker et al. 2010, Batistic 2012, Lin and Conibear 2015, Li et al. 2017, Liu et al. 2021). Many studies, primarily conducted in animals, indicate that S-acylation is important in regulating the trafficking, localization, activity, stability, and interactions of proteins. Therefore, protein S-acylation likely plays crucial roles in a wide range of biological processes, such as cellular signaling, metabolism, and pathogenesis (Li et al. 2017, Turnbull and Hemsley 2017).

In mammals, the functions of protein S-acylation and their regulatory mechanisms have been extensively studied. Many proteins involved in the regulation of growth, development, and a wide range of diseases are known to be S-acylated (Chen et al. 2017, Zhang et al. 2021). The enzymes involved in S-acylation and de-S-acylation processes have also been well studied (Li et al. 2017). In contrast, research on protein S-acylation in plants lags, and the regulation mechanism of protein S-acylation in plants is still not clear. Nevertheless, an increasing number of proteins have been identified as S-acylated through biochemical or genetic analysis, such as heterotrimeric G proteins, specific cellular signal transduction components (Adjobo-Hermans et al. 2006, Zeng et al. 2007), small GTPases AtROP6/9/10 (Lavy et al. 2002, Zheng et al. 2002, Sorek et al. 2017), proteins involved in Ca^2+^ signaling (e.g., Calcineurin B-Like proteins AtCBL1/2/3/6 in *Arabidopsis* and the calcium dependent protein kinase OsCPK2 in rice) (Martín and Busconi 2000, Batistic et al. 2008, Batistič et al. 2012), immune response-related proteins [e.g., Flagellin-Sensitive 2 (FLS2), extracellular ATP receptor P2K1, AvrPphB Susceptible1 (PBS1), PBS1-Like 1 (PBL1), PBS1-Like 19 (PBL19), and R5L1] (Kim et al. 2005, Hemsley et al. 2013, Qi et al. 2014, Chen et al. 2021, Liu et al. 2021, Gao et al., 2022), and cellulose synthesis-related proteins CESA1/4/6/7/8 (Kumar et al. 2016, 2022).

PATs are integral membrane proteins with a highly conserved Asp-His-His-Cys (DHHC) motif, responsible for adding fatty acyl groups to cysteine residues in substrate proteins (Roth et al. 2002, Zheng et al. 2019). Twenty-four PATs, named AtPAT1-AtPAT24, are predicted to be encoded by the *Arabidopsis* genome (Batistič et al. 2012). So far, only a few of the PAT-substrate pairs have been identified and functionally characterized: e.g., PAT4 S-acylates ROP2 at the plasma membrane and regulates root growth (Wan et al. 2017); PAT10 S-acylates CBL2/3/6/10 at the tonoplast membrane to regulate salinity tolerance (Zhou et al. 2013, Chai et al. 2020); and PAT13/14 possibly S-acylate Nitric Oxide Associated 1 (NOA1) within chloroplasts to influence leaf senescence (Lai et al. 2015); PAT13- and PAT16-SLacylates the NBSLLRR protein R5L1 to mediate its membrane localization to activate the plant defense response (Gao et al., 2022); PAT24 interacts with and S-acylates cellulose synthase complexes (CSCs) to influence cellulose biosynthesis in plant leaves (Zhang et al., 2022, Lampugnani et al., 2024). Our recent study also discovered that PAT5 and 9 S-acylated the extracellular ATP receptor P2K1 at the plasma membrane to regulate the immune response in *Arabidopsis* (Chen et al. 2021). Interestingly, P2K1 was shown to directly phosphorylate and stimulate S-acylation activity of PAT5 and PAT9. The activated PAT5 and PAT9, in turn, S-acylated P2K1 to promote P2K1 degradation to dampen immune responses (Chen et al. 2021). This example clearly illustrates the intricate relationship between PATs and their substrates. Since the number of potential S-acylated proteins in *Arabidopsis* far exceeds the number of PATs, each PAT likely S-acylates multiple substrates. PAT5 and PAT9 likely also S-acylate other proteins in addition to P2K1.

S-acylproteomic methods were developed first in yeast, and then in mammals and plants to identify S-acylated proteins (Wang and Yang 2021). Due to the lack of S-acylation specific antibodies, S-acylproteomic studies were conducted by indirect S-acylation enrichment methods, coupled with liquid chromatography-mass spectrometry (LC-MS). Three major quantitative S-acylproteomic approaches were developed to identify S-acylated proteins (Wang and Yang 2021): metabolic labeling with a palmitic acid analogue, followed by click chemistry (MLCC) (Martin and Cravatt 2009, Sobocińska et al. 2017, Won and Martin 2018), acyl resin-assisted capture (acyl-RAC) (Forrester et al. 2011, Edmonds et al. 2017, Kumar et al. 2022), and acyl-biotinyl exchange (ABE) (Hemsley et al. 2013, Edmonds et al. 2017, Collins et al. 2017, Zhou et al. 2019, 2020). MLCC is a metabolic-dependent method and has been rarely applied to analyze S-acylation-sites (Boyle et al 2016, Wang and Yang 2021)The ABE and acyl-RAC methods do not require metabolic labeling, making them universally applicable. Furthermore, the ABE method can be coupled with both protein-level and peptide-level (also termed as site-specific) S-acylation enrichment approaches to globally identify S-acylated proteins as well as their S-acylation sites (Collins et al. 2017). Due to these advantages, the ABE method has become the most widely used approach.

In plants, two major S-acylproteomic studies were conducted using the ABE-based protein-level enrichment method (Hemsley et al. 2013, Srivastava et al. 2016). About 600 and 450 putative S-acylated proteins were identified in *Arabidopsis* root cell suspension culture (Hemsley et al. 2013) and poplar cell suspension culture (Srivastava et al. 2016), respectively. However, no exact S-acylation sites were identified directly from these studies. A recent S-acylproteomic study was performed in different tissues of *Arabidopsis* using the acyl-RAC-based peptide-level enrichment approach. Over 5000 S-acylated cysteine sites, matching 2643 proteins, were identified (Kumar et al. 2022). The variations observed among these reports imply significant differences in the S-acylproteomes across different tissue types and under different conditions. These variations may also be caused by potential methodological biases.

In the present study, our main objectives were to obtain a comprehensive *Arabidopsis* S-acylproteome, and to identify potential substrates for PAT5 and PAT9. Given that PATs primarily S-acylate membrane proteins or membrane-associated proteins, as well as our laboratory’s interest in plasma membrane-associated receptor proteins, we focused on S-acylation of membrane proteins. For this purpose, S-acylated cysteines from isolated membrane proteins were first enriched via ABE method at both protein (ABE-protein) and peptide (ABE-peptide) levels. The enriched samples were then analyzed using label-free LC-MS. Detailed analyses of the data showed that the two methods were complementary in identifying S-acylated proteins. Together they revealed over 2000 putative S-acylated proteins, many of which were not previously revealed. Additionally, over 100 potential substrates were identified for both PAT5 and PAT9. Thus, our study not only expanded the *Arabidopsis* S-acylproteome, but also identified a variety of potential substrates for the two important plant immunity-related protein S-acyltransferases, PAT5 and PAT9. These results provide an excellent resource for further functional analyses of S-acylated proteins in plants.

## RESULTS

### Application of the ABE-protein and ABE-peptide Methods to study the *Arabidopsis* S-acylproteome

To obtain a global, comprehensive picture of the S-acylated proteins (especially for the membrane proteins) and S-acylation sites in *Arabidopsis*, both the protein-level and peptide-level enrichment approaches were used with subsequent LC-MS analysis (Figure 1). Tandem mass (MS/MS) spectra were searched against the *Arabidopsis thaliana* proteome in the UniProt database (https://www.uniprot.org/) using PEAKS Studio. From the ABE-protein data, a total of 57,098 unique peptides, matching 5,530 *Arabidopsis* proteins, were identified at an FDR of 1% from all six genotypes (Col-0, *pat5*, *pat9*, *pat5*/*pat9*, c-*pat5* and c-*pat9*). From the ABE-peptide data, a total of 9,482 peptides, matching 1,537 *Arabidopsis* proteins, were identified. To compare the two enrichment methods, Col-0 was used to analyze cysteine modifications (Figure 2A, Tables S1, S2). In the ABE-protein data of Col-0, only 5,631 (11%) out of 50,614 detected unique peptides contained at least one Cys: 5,218 with N-ethylmaleimide (NEM), 257 with free Cys, 120 with CAM, and 36 with 2,2′-dithiodipyridine (DTDP) modification, with a proportion of 92.7%, 4.6%, 2.1%, and 0.6%, respectively, in the Cys-containing peptides. The ABE-peptide data of Col-0 consisted of a total of 6,112 unique peptides, including 5,075 (83%) Cys-containing peptides. Among the Cys-containing peptides, 344 were identified with NEM, 743 with free Cys, 3,984 with CAM, and 4 with DTDP, with a proportion of 6.8%, 14.6%, 78.5%, and 0.1%, respectively. Although the total number of peptides/proteins revealed by the ABE-peptide method was smaller, this approach identified significantly more peptides with CAM modifications, which are most likely to be S-acylated *in vivo*. Since non-S-acylated cysteines may have been labeled by IAM if they were not effectively blocked by NEM or DTDP, and on the other hand, some S-acylation sites may have become free cysteine residues due to insufficient IAM alkylation. It is critical to compare the proteomics data between the enriched samples (+Hyd) and control samples (-Hyd) quantitatively. This analysis will help unveil potentially real S-acylated proteins while minimizing any artifacts.

**Figure 1.**
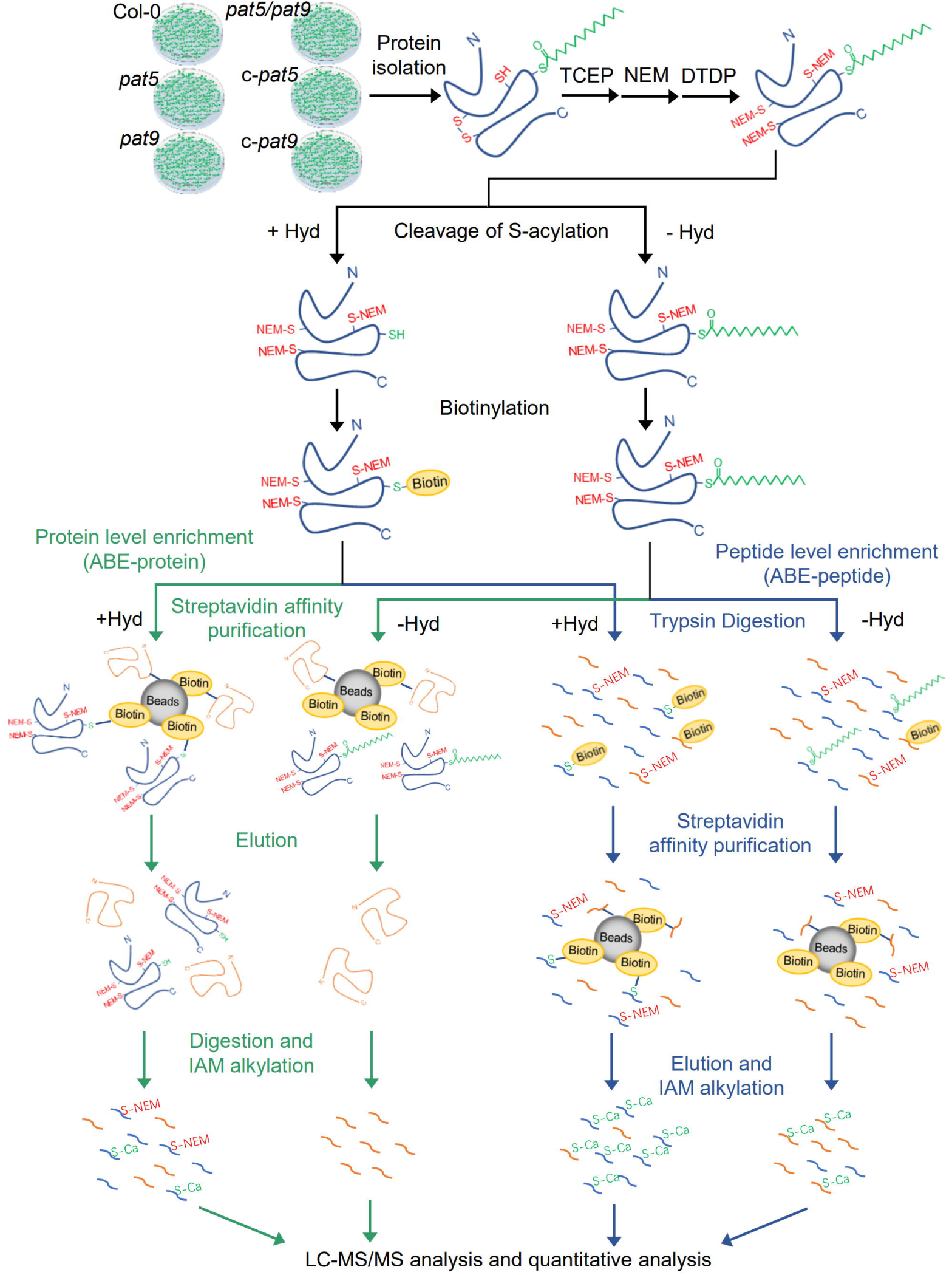
Schematic illustration of S-acyl proteome analysis describing both protein-level (ABE-protein) and peptide-level (ABE-peptide) enrichment methods to identify S-acylated proteins and substrates of PAT5/PAT9 in *Arabidopsis*. 14-day-old seedlings of wild type Col-0, mutant *pat5*, *pat9*, *pat5/9*, and complemented lines c-*pat5*, c-*pat9* were prepared and membrane proteins were isolated and enriched. Proteins were reduced by TCEP, and free cysteines were blocked by N-ethylmaleimide (NEM) and 2,2′-dithiodipyridine (DTDP). One sample was subsequently equally divided to two, named +Hyd (enriched sample) and -Hyd (control sample), and underwent acyl-biotin exchange (ABE) procedure: in +HAM sample, S-acyl groups were specifically removed by hydroxylamine (Hyd) from cysteines and newly released cysteines were biotinylated by HPDP-Biotin; in -Hyd sample, Hyd was absent and biotinylated sites were background noise. Two methods were used to enrich the biotinylated cysteine residues: 1), protein-level enrichment, proteins with biotinylated cysteines were captured and purified by streptavidin agarose resin. Enriched proteins were then trypsin-digested; 2), peptide-level enrichment, trypsin digestion was conducted before the resin capture. Digested peptides underwent streptavidin affinity purification procedures. After the S-acylation-sites enrichment, peptides were alkylated by iodoacetamide (IAM) and identified by label-free mass spectrometry. SH, thiol groups of cysteine residues; S, cysteine sulfur; Ca, carbamidomethylation by IAM.

**Figure 2.**
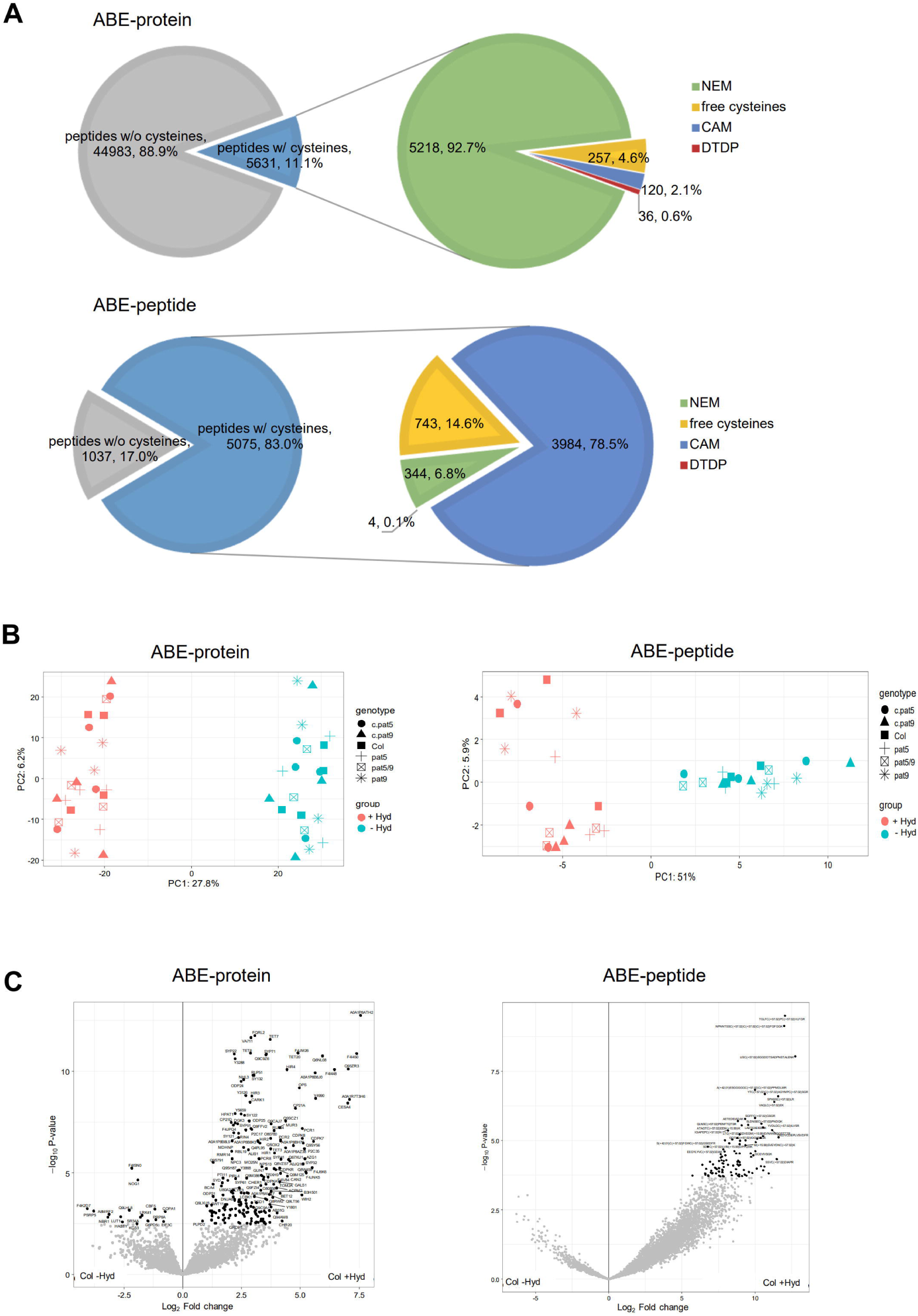
Overview of the S-acyl-proteomes by two enrichment methods. (A) Pie charts showing the distribution of detected peptides with or without cysteines (left) and modifications on cysteines (right) in ABE-protein (top) and ABE-peptide (bottom) S-acylproteomes. Numbers and percentages in the pie chart indicate the peptide amounts and proportions in total detected peptides. NEM, N-ethylmaleimide on cysteines; DTDP, 2,2′-dithiodipyridine on cysteines; CAM, carbamidomethylation on cysteines, candidate S-acylation sites; free cysteines, free cysteines without any modifications. (B) PCA analysis for ABE-protein (left) and ABE-peptide (right) S-acylproteome from seedlings of Col-0, *pat5*, *pat9*, *pat5/9*, and c-*pat5*, c-*pat9*. Samples treated with (+Hyd) or without hydroxylamine (-Hyd) were S-acylation-enriched samples or background controls. (C) Volcano plots showing the enrichment of proteins/peptides in Col +Hyd (Col-0 with Hyd treatment) compared to Col -Hyd (Col-0 without Hyd treatment) in both ABE-protein (left) and ABE-peptide (right) S-acylproteomes.

Principal component analysis (PCA) (Figure 2B) of both ABE-protein and ABE-peptide S-acylproteome data showed a good separation between the S-acylation enriched samples (+Hyd) and the control samples (-Hyd). However, the wild-type Col-0 and mutants were not well separated, suggesting that the difference between the S-acylproteomes in the wild type and mutants may be masked by the big overlap among genotypes and variance among replicates. Another possible reason is that PAT5 and PAT9 are only responsible for S-acylation of a relatively small number of proteins. Volcano plots (Figure 2C, Figure S1) from both ABE-protein and ABE-peptide S-acylproteomes also showed a significant difference between the +Hyd samples and the -Hyd controls. Therefore, we will compare the +Hyd samples and the -Hyd controls to identify potential S-acylated proteins in the next section.

### Identification of Putative S-acylated Proteins using the ABE-protein and ABE-peptide Methods

From the aforementioned data, a large number of proteins/peptides enriched in the +Hyd samples were revealed in Col-0, as well as in the other genotypes (Figure 2C, Tables S3, S4). To identify possibly real S-acylated proteins/peptides, the +Hyd samples and the -Hyd controls were compared: Proteins/peptides with a fold change (FC) > 2 and FDR < 0.05 were considered significantly enriched, thereby designating them as putative S-acylated proteins/peptides. Using these criteria, 393 to 469 S-acylated proteins (Figure 3A) were identified in the ABE-protein data for each genotype. Similarly, 1054 to 1657 S-acylated peptides were identified in the ABE-peptide data (matching 755 to 908 proteins) for each genotype. Thus, the ABE-peptide method revealed more putative S-acylated proteins than the ABE-protein method.

**Figure 3.**
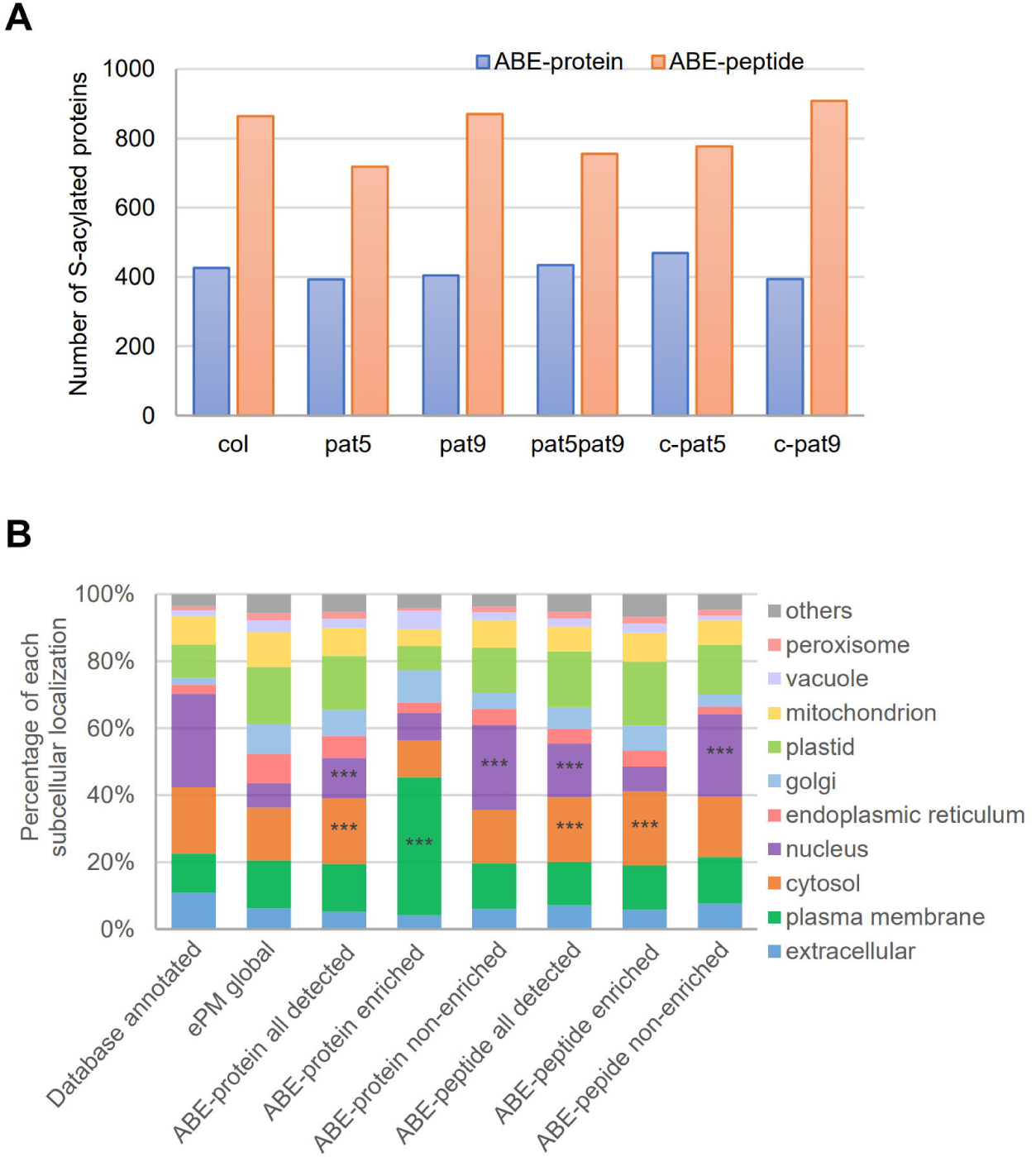
S-acylated proteins identified by the S-acyl-proteomes. (A) Number of S-acylation enriched proteins by two enrichment methods from seedlings of six genotypes. (B) Subcellular distribution of protein profiles from the database annotated proteins, global proteome of the enriched plasma membrane (ePM) proteins, and S-acyl proteomes (ABE-protein and ABE-peptide). Protein subcellular localization was predicted using SUBA4. Compared to the ePM global proteome, the subcellular enrichment of the protein categories in the S-acyl-proteomes was tested using the hyper-geometric distribution. Overrepresented localizations (p < 0.001) are indicated with ***.

To further illustrate the differences between the two methods, intensity distributions and fold enrichment of S-acylated proteins in Col-0 were analyzed. In the ABE-protein data, log2 transformed intensities of S-acylated proteins identified in the +Hyd and -Hyd samples displayed a similar gourd shape in the violin plot with a FC 6.5 (log2 2.7) at the median line (Figure S2). In the ABE-peptide data, log2 intensities of S-acylated proteins presented a more rounded gourd shape in the +Hyd samples and a half gourd in the -Hyd samples with a FC 71.0 (log2 6.2) at the median line. The formation of the lower half gourd in the plot may be due to the imputation of non-detected intensities. The absence of the upper-half gourd in the -Hyd samples indicated the high purity of the enrichment via the ABE-peptide method. Surprisingly, only 62 proteins (17% of proteins identified in ABE-protein and 8% of proteins identified in ABE-peptide) were found S-acylated in Col-0 in both S-acylproteomes (Figure S3), indicating the high complementarity of the two approaches in identifying S-acylated proteins.

### Comparison of our *Arabidopsis* S-acylproteomes with the reported S-acylproteomes

To evaluate the quality of our data, we compared our Col-0 S-acylproteomes with two previously reported *Arabidopsis* S-acylproteomes. The comparison revealed both similarities and differences (Figure S4). Approximately 77% and 30% S-acylated proteins identified from the ABE-peptide and ABE-protein approach, respectively, were previously reported in the acyl-RAC based peptide-level S-acylproteomics study of *Arabidopsis* seedlings (Kumar et al. 2022). In contrast, about 18% and 30% of the S-acylated proteins identified from our two approaches, respectively, were previously reported in the ABE based protein-level S-acylproteomics study of *Arabidopsis* root cell suspension (Hemsley et al. 2013). Notably, 380 proteins identified in our study were not previously reported as S-acylated proteins, suggesting that they may represent novel S-acylated proteins (Table S5). The differences between these various studies may be attributed to the use of differing methods and biological materials, and the specific focus of our study on membrane proteins. However, the same enrichment method (protein-level or peptide-level) yielded more common S-acylated proteins. These findings indicate the high quality of our data and underscore a significant difference between the two approaches in identifying S-acylated proteins.

### Subcellular localizations of the identified S-acylated proteins

To evaluate the effectiveness of our methods in identifying S-acylated membrane proteins, subcellular localizations of the identified S-acylated proteins were predicted using SUBA4. The overrepresentation of subcellular localizations was evaluated against the global proteome of the starting proteins for S-acylation enrichment, i.e., enriched PM proteins (ePM) (Figure 3B and Figure S5). Compared with all the annotated proteins from the TAIR database, the percentages of the nuclear and cytosolic proteins in the ePM proteome were decreased, and the percentages of the proteins localized in the membrane-containing organelles (vacuole, mitochondrion, plasmid, Golgi, ER, and PM) were increased, indicating a successful enrichment of membrane proteins during the protein isolation procedures. In the ABE-protein S-acylproteome, PM proteins were overrepresented in the enriched proteins (enriched proteins 41% vs. ePM 14%), while the nuclear proteins were overrepresented in the non-enriched proteins (non-enriched proteins 25% vs. ePM 7%). On the contrary, the ABE-peptide S-acylproteome showed an overrepresentation of cytosolic proteins (enriched proteins 22% vs. ePM 16%) (Figure S5). Therefore, the ABE-protein method is more effective in identifying PM-associated S-acylated proteins.

### Functional Annotation of the Identified S-acylated Proteins

To understand the potential biological functions associated with S-acylation in *Arabidopsis*, we performed gene ontology (GO) enrichment analysis using the putative S-acylated proteins from Col-0 (Figure 4). The S-acylated proteins identified using the ABE-protein approach were enriched in the following GO biological processes: protein phosphorylation, vesicle fusion, vesicle-mediated protein transport, cellular signal transduction, and immune responses (Figure 4A). Correspondingly, GO terms of protein kinase activity, calmodulin-dependent protein kinase activity, SNAP receptor activity, S-acyltransferase activity, etc., were enriched in the molecular function category. Cellular components involved in endomembrane system, vesicle/protein trafficking, and cell wall were also enriched. These findings agree with the reported S-acylproteomic studies and functional analyses of S-acylated proteins in plants and other organisms (Hemsley et al. 2013, Srivastava et al. 2016, Collins et al. 2017). Interestingly, except that the vesicle-mediated transport (Golgi to PM) is consistent with the GO term in the ABE-protein acylproteome, many top enriched GO terms in the ABE-peptide acylproteome (Figure 4B) were related to organ development, primary biosynthesis, and metabolic processes. The GO result from the ABE-peptide data was similar to that from the acyl-RAC-based peptide-level enrichment S-acylproteome in *Arabidopsis* (Kumar et al. 2022). These findings further demonstrate that the two approaches preferentially captured different groups of proteins and are therefore complementary in identifying S-acylated proteins from an organism.

**Figure 4.**
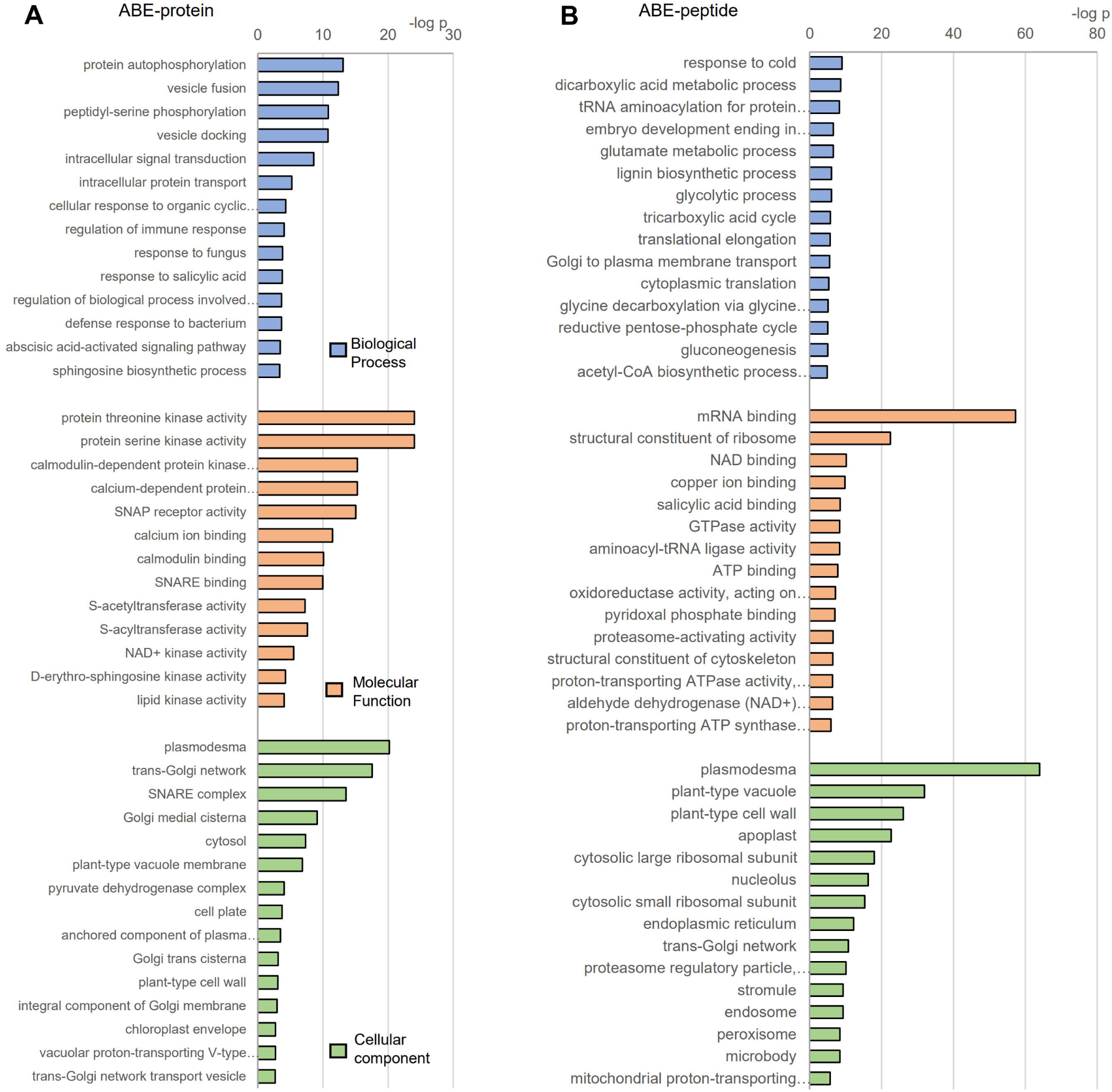
GO enrichment analysis of S-acylated proteins identified using ABE-protein (A) and ABE-peptide methods from Col-0 (B). Representative enriched GO terms identified by *PANTHER* for each cluster in the category of biological processes, molecular functions, and cellular components are shown.

Protein biological functions are often mediated by protein-protein interactions (PPIs) and S-acylation may impact PPIs. Therefore, we further investigated possible PPIs for the S-acylated proteins (from the ABE-protein data) employing the STRING database. Many PPIs were revealed with high confidence as shown in Figure S6. Four big interaction clusters, potentially involved in vesicle-mediated transport, protein biosynthesis, phosphorus metabolic process, and cell surface receptor signaling (part of GO term of response to stimulus), were identified. Consistent with the GO analysis and previous studies (Greaves et al. 2010, Hemsley et al. 2013, Collins et al. 2017, Zhou et al. 2019), many interacting soluble N-ethylmaleimide-sensitive factor-activating protein receptors (SNAREs), responsible for vesicle trafficking and vesicle-mediated protein transport, were predicted to be S-acylated. Notably, over 10 Leucine-rich repeat protein kinase family proteins, such as FLS2, Guard cell Hydrogen peroxide-Resistant 1 (GHR1), STRUBBELIG-receptor family (SRF) proteins, and leucine-rich repeat protein 1 (LRR1), subjected to the GO term of cell surface receptor signaling, were identified to be S-acylated. These results suggest that protein S-acylation may impact a broad range of biological functions through regulating PPIs.

### Motif Analysis of S-acylation Sites

To reveal any consensus motifs associated with S-acylation, we examined the flanking sequences of S-acylated cysteines from the ABE-peptide data with the pLogo algorithm. Since S-acylation can impact protein subcellular localizations, the S-acylated proteins in each subcellular localization category were individually analyzed. The analysis showed that many residues were overrepresented (Figure 5), e.g., valine (V) at −1, alanine (A) at +1 position of the S-acylation sites from the cytosol localized S-acylated proteins; lysine (K) at −7 position from the ER localized S-acylated proteins; threonine (T) at −1 and proline (P) at +1 position from the extracellular S-acylated proteins, etc. Interestingly, except the PM and extracellular category, cysteines around the S-acylation sites were under-represented. In the PM proteins, cysteines appeared overrepresented (although not statistically significant) in the ±1 position, indicating that double or triple S-acylation in PM proteins may be common. Intensive S-acylation modifications, perhaps in conjunction with other lipid modifications, can increase the hydrophobic nature and binding affinity of proteins to the PM, as shown in some PM proteins, e.g., triple Cys in C-terminal domain of RIN4 (Kim et al. 2005), several double/triple Cys in variable region 2 (VR2) of CESA7 (Kumar et al. 2016), and a combination of S-acylation with myristoylation/prenylation on heterotrimeric G proteins (Adjobo-Hermans et al. 2006, Zeng et al. 2007). In addition, basic amino acids, such as lysine (K) and arginine (R), are underrepresented at the −1 position of the S-acylation sites, indicating that modifications on K/R residues, such as acetylation, methylation, or ubiquitination is not likely to occur next to an S-acylation site.

**Figure 5.**
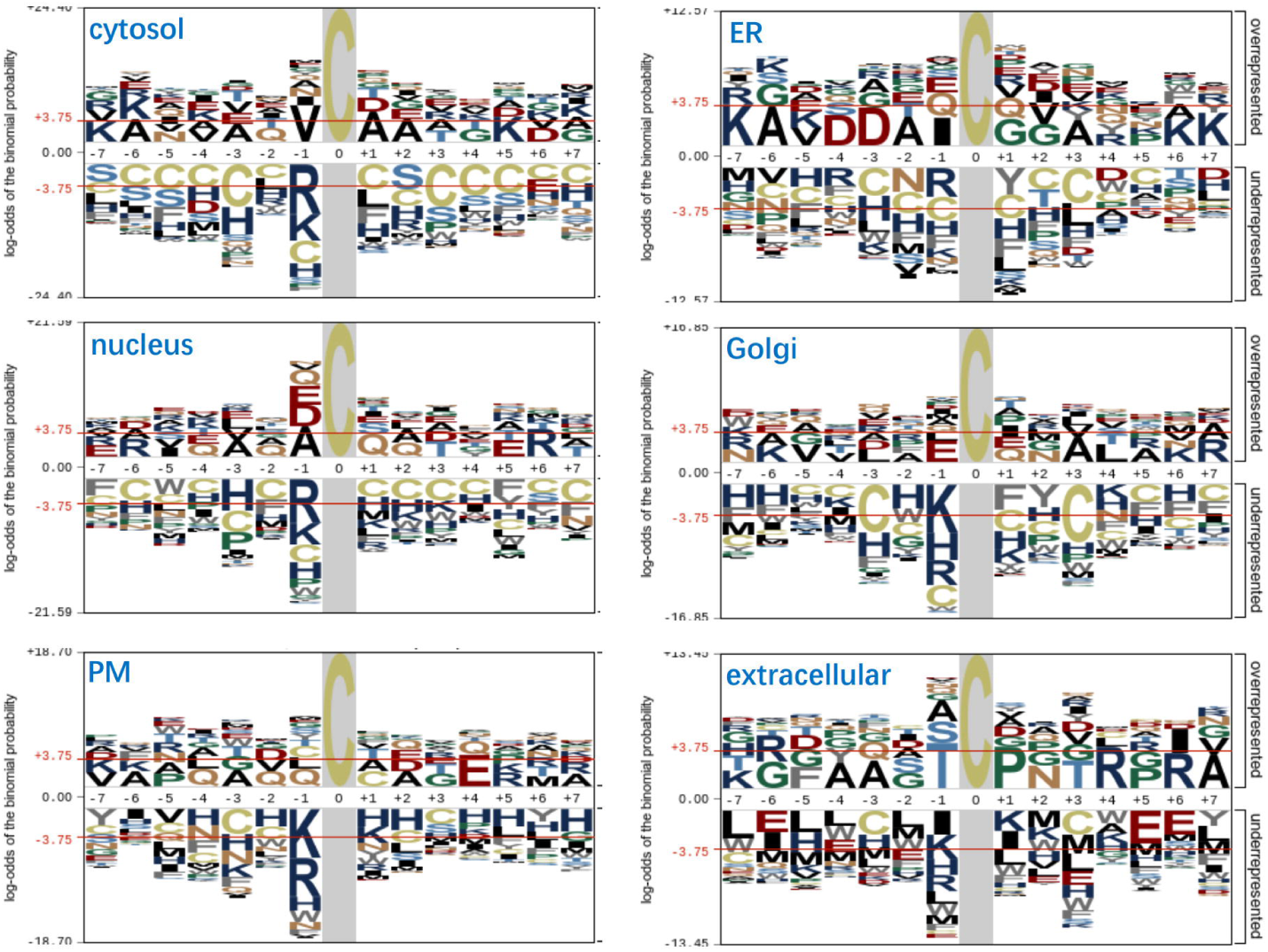
Motif analysis of S-acylated peptides identified from the ABE-peptide method. Peptides from proteins localized in each subcellular component were analyzed separately. Analysis was performed with pLogo using a 15 amino acid long peptide sequence containing the central cysteine and 7 residues on either side. The *Arabidopsis* proteome was used as background. The red horizontal lines on the plogo plots denote p < 0.05 thresholds. Residues that exceed the threshold lines are over-represented (above 0 line) or under-represented (below 0 line). ER, endoplasmic reticulum; PM, plasma membrane.

### New Findings for Previously Reported S-acylated Proteins

Our study also revealed additional S-acylation information for several previously reported S-acylated proteins. Our study also obtained a high coverage of proteins with multi-S-acylations. For example, RIN4, a member of the R protein complex and targeted by at least three bacterial virulence factors, needs one or more of the three consecutive cysteines in its C-terminus for protein stabilization and localization on the PM, and these cysteines are likely S-acylated or prenylated (Kim et al. 2005). Indeed, both the ABE-protein and ABE-peptide data showed that RIN4 was S-acylated and S-acylation was detected at the following cysteine residues: C(203), C(204), C(205) from a number of peptides (Figure 6A). The variations in S-acylation at these three cysteines may have differential effects on the localization of RIN4 and potentially impact its function.

**Figure 6.**
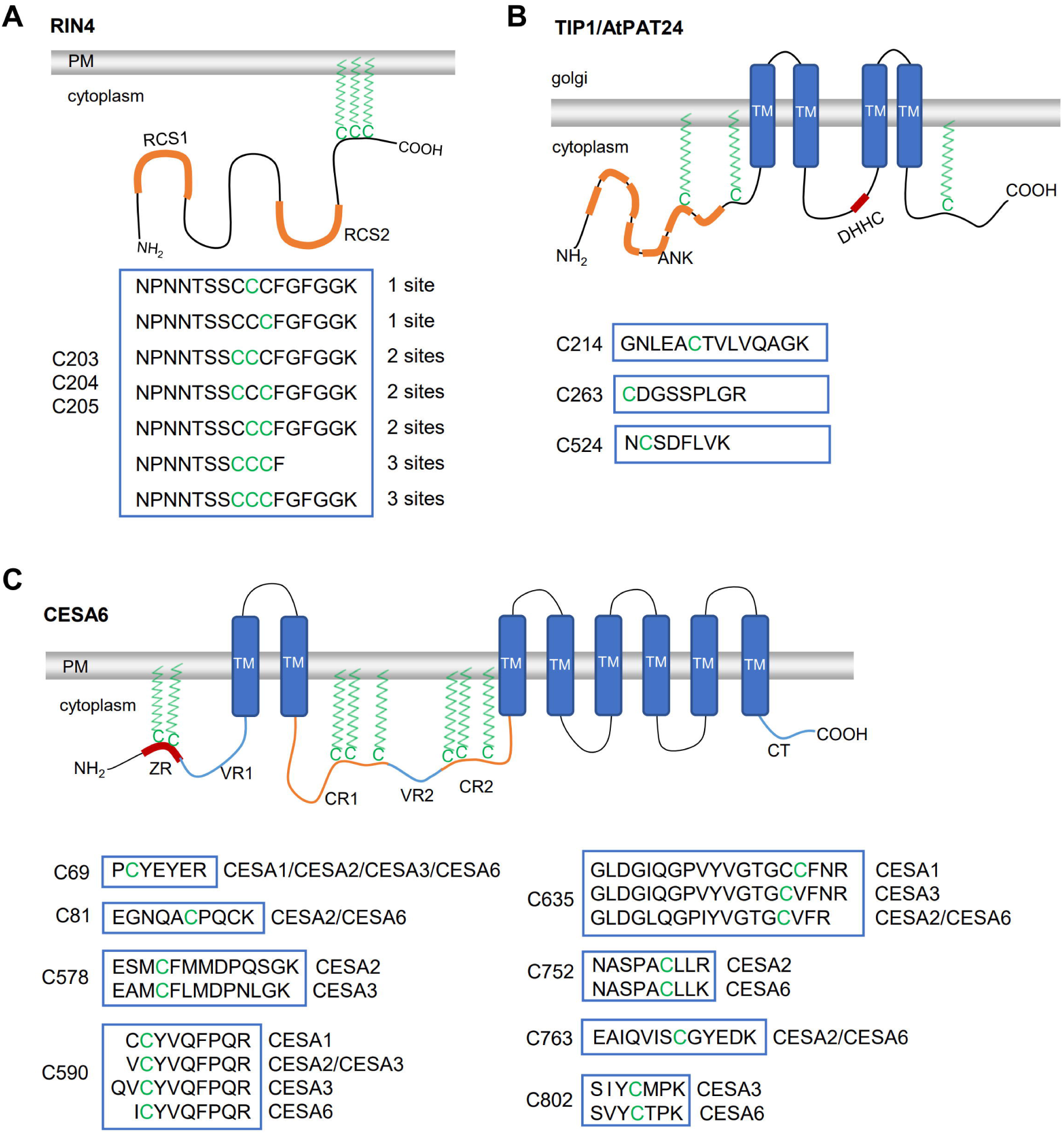
Schematic of identified S-acylation sites on representative, known S-acylated proteins. The S-acylation sites are indicated on the protein architectures with functional domains. The enriched peptide sequence windows detected in the data are indicated and cysteines with S-acylation are highlighted in green. (A) Non-transmembrane protein RIN4. PM, plasma membrane. RCS1, AvrRpt2 cleavage site 1; RCS1, AvrRpt2 cleavage site 2. (B) Golgi localized transmembrane protein TIP1. TM, transmembrane domain; ANK, ankyrin repeat domain; DHHC, DHHC cysteine-rich domain. (C) Plasma membrane-localized transmembrane protein family CESAs. TM, transmembrane domain; ZR, Zinc RING type finger; VR1, variable region 1; VR2, variable region 2; CR1, conserved region 1; CR2, conserved region 2; CT, carboxy terminus.

In addition to S-acylating other proteins, PATs themselves can be S-acylated at the DHHC motif and outside this motif as revealed in animals (Mitchell et al. 2010, Zmuda and Chamberlain 2020). So far, no S-acylation sites have been identified for any plant PATs. Our current study revealed three S-acylation sites on AtPAT24: C214 (in the ankyrin repeat domain, which is thought to be responsible for the substrate specificity) (Lemonidis et al. 2015), C263, and C524 (C-terminal) (Figure 6B). AtPAT24, also called Tip Growth Defective 1 (TIP1), whose loss-of-function mutant shows pleiotropic defects in root hair growth and other aspects, is one of the few well-characterized plant PATs (Hemsley et al. 2005). These three S-acylation sites might be important for the proper function of PAT24 in substrate specificity, catalytic activity, and/or protein localization.

CESA proteins are also well-studied S-acylated proteins. They are essential for cellulose synthesis in both the primary (CESA1/3/6) and secondary (CESA4/7/8) cell wall (Polko and Kieber 2019). The S-acylation of Cys in the variable region 2 (VR2) and at the carboxy terminus (CT) of CESA7 is required for proper protein function by regulating protein trafficking from the Golgi to the PM (Kumar et al. 2016, 2022). Our current study found that two Cys in the Zinc RING type finger (ZR), three Cys in the conserved region 1 (CR1), and three Cys in the CR2, were S-acylated in CESA6 and CESA1/2/3 (Figure 6C). Most of these sites (C69, C578, C590, C635, C763, C802) were also identified by Kumar *et al*. (2022), suggesting that these cysteine residues were very likely S-acylated in plants. Two previously unknown S-acylation sites, C81 and C752, within CESA6, were also identified in our study. The C69 is highly conserved among CESA1/2/3/6, so that all four CESAs might be S-acylated at this site. In addition, S-acylated C590 and C635 in CESA6 were also detected in three other CESAs, suggesting a conserved function for these sites across multiple CESA proteins. The S-acylation sites revealed in the ZR region of CESA6 suggest that CESA6 may function differently from CESA7, in which S-acylation was not predicted to occur in the ZR region due to its role in zinc binding (Kumar et al. 2016).

### Analysis of S-acylation of RLKs

Previous studies showed that S-acylation is required for proper function of receptor-like kinases (RLKs)/receptor-like cytoplasmic kinases (RLCKs) on the plasma membrane, including pathogen responses and downstream cellular signaling (Hemsley et al. 2013, Qi et al. 2014, Hurst et al., 2019, 2023, Chen et al. 2021). Consistent with this, S-acylated RLKs were found overrepresented in the GO and PPI analyses. Overall, 63 RLKs (including 22 RLCKs) from 27 gene families, potentially involved in various biological functions, were detected in our study with different enrichment levels in Col-0 (Figure S7). Forty-five potential S-acylation sites from 22 RLK/RLCK proteins were identified using the ABE-peptide method.

To reveal any possible S-acylation distribution patterns in proteins localized in each cellular component and RLKs, the relative positions of S-acylation sites were calculated along the protein lengths and depicted in a violin plot (Figure 7A). Interestingly, the distribution of S-acylation sites in the proteins from the different organelles showed differing patterns. Except proteins in the mitochondrion and vacuole, S-acylation sites were generally distributed through the entire length of the protein, with a higher frequency in the 0.3 - 0.6 and C-terminal region of the proteins. The proteins from mitochondria showed intensive S-acylation at both ends, while the proteins in vacuoles showed an opposite pattern. Compared with the other regions in the RLK proteins, less S-acylation was detected in the 0.5 - 0.8 region (Figure 7A).

**Figure 7.**
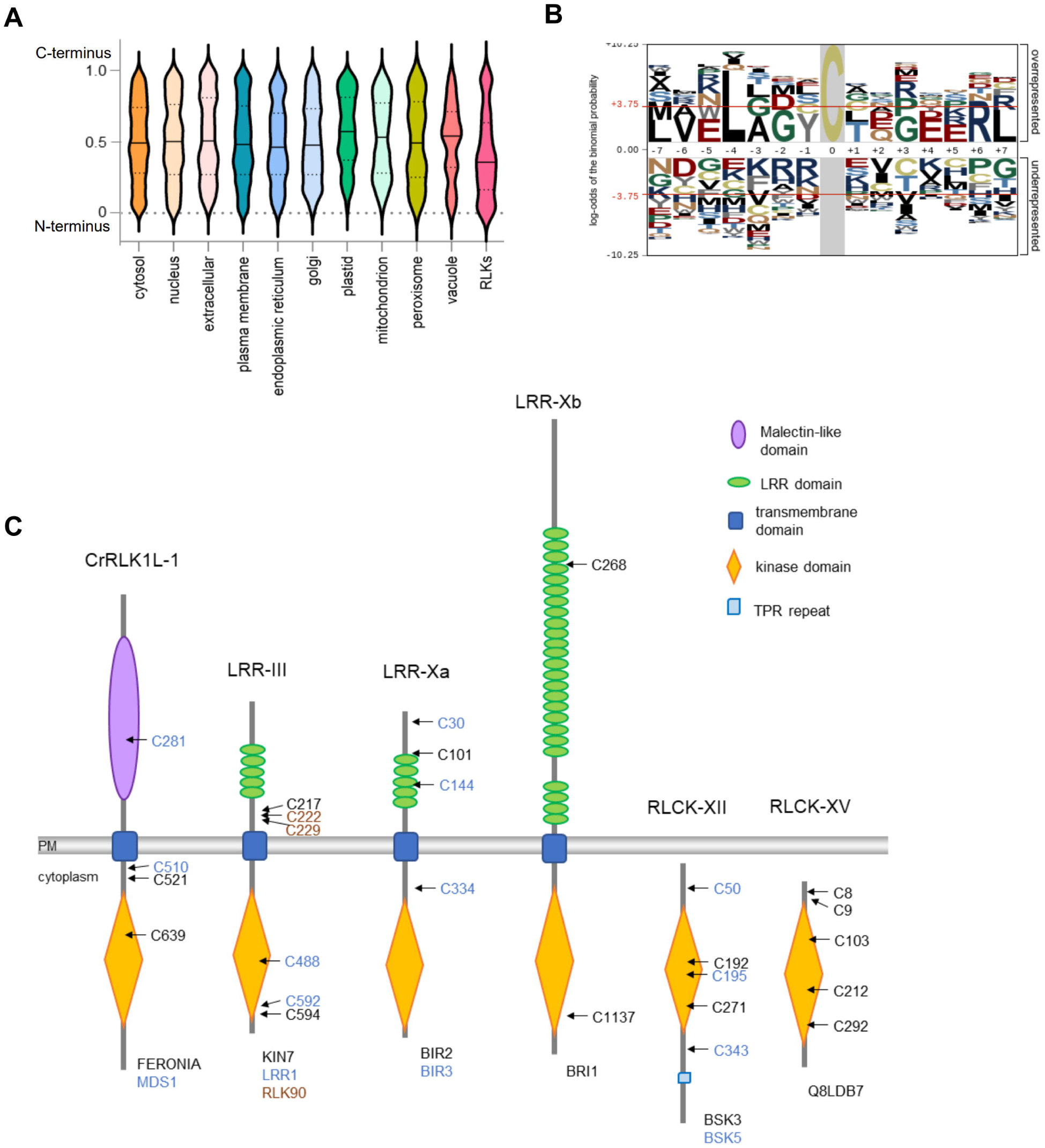
Analysis of S-acylation in RLKs. (A) Violin plot of position distributions of the S-acylation sites identified by the ABE-peptide method relative to protein length. Peptides from proteins localized in each subcellular component were separately analyzed. (B) Motif analysis of S-acylated peptides from RLKs. Analysis was performed with plogo using a 15 amino acid long peptide sequence containing the central cysteine and 7 residues on either side. The *Arabidopsis* proteome was used as background. The red horizontal lines on the plogo plots denote p < 0.05 thresholds. (C) Representative schematics of protein architecture showing the S-acylation sites for RLK proteins. Protein families and S-acylated protein names are indicated below the protein architectures. Functional protein domains are represented with different shapes and colors. S-acylation sites for each protein are shown with distinguishable colors.

Similar to the PM proteins, the RLKs showed an overrepresentation of Cys at the ±1 position relative to their S-acylation sites different from the proteins in other cellular components (Figure 7B). Furthermore, leucine (Leu) was also overrepresented at the −4 to +1 position of an S-acylation site in these RLKs. We initially speculated that the enrichment of Leu was due to the leucine-rich feature in the LRR-rich receptor kinase families. However, this enrichment was commonly observed across all RLK families, including RLCKs, suggesting that the overrepresentation of Leu at the −4 to +1 positions may hold biological significance for this group of proteins.

As shown in Figure 7C, S-acylation was detected in multiple domains of RLKs/RLCKs, including the kinase domain, LRR domain in the LRR-RLK families, the malectin-like domain and the juxta-transmembrane domain in CrRLK1L family. S-acylation in different domains may lead to different impacts on protein function. S-acylation sites on the RLCKs, such as Brassinosteroid Signaling Kinase 3/5 (BSK3/5), might be crucial to protein localization on the PM, similar to the reported S-acylated RLCK SCHENGEN1 (SGN1) (Alassimone et al. 2016). But S-acylation on RLKs is not likely to determine their general targeting to the membrane due to the presence of the transmembrane domain,. S-acylation in these RLKs may impact their activity, stability, and trafficking through the vesicle-mediated protein sorting system.

### Identification of Potential Substrates of PAT5 and PAT9

Quantitative S-acylproteomics has been successfully employed to discover PAT-substrate pairs by comparing the differences of S-acylproteomes in WT and PAT-deficient cells or tissues in yeast, mammalian systems, and plants (Roth et al. 2006, Hemsley et al. 2013, Shen et al. 2017, Gorinski et al. 2020). To identify potential substrates of PAT5 and PAT9, S-acylated proteins identified from wild-type Col-0, single mutants *pat5* and *pat9*, double mutant *pat5/9*, and the complemented lines of *pat5* (c-*pat5*) and *pat9* (c-*pat9*) were compared. Potential PAT5 and PAT9 substrates should be enriched in the +Hyd samples (compared to -Hyd) of Col-0 and the complemented lines, and at the same time show lower or no enrichment in the single and double mutants. To simplify the analysis, putative S-acylated proteins identified in Col-0 and the complementary lines, but absent in the corresponding mutants, were considered the candidate substrates for these PATs.

Using the above criterion, 20 and 11 proteins were identified to be potential substrates of PAT5 and PAT9, respectively, from the ABE-protein S-acylproteome (Figure 8A). These proteins belong to diverse GO Biological Processes (Figure 8B), such as regulation of defense responses (CDPKS, PBL40), cellular signal transduction (F4JR41, RAC10), and cell wall biogenesis (PMTI, F4IVP3). Considering the PM localization of PAT5 and PAT9, those candidate proteins localized on PM are more likely the real substrates. Therefore, three PM localized proteins, A0A178UFH0, CI111, and MUC70, were likely true substrates for both PAT5 and PAT9.

**Figure 8.**
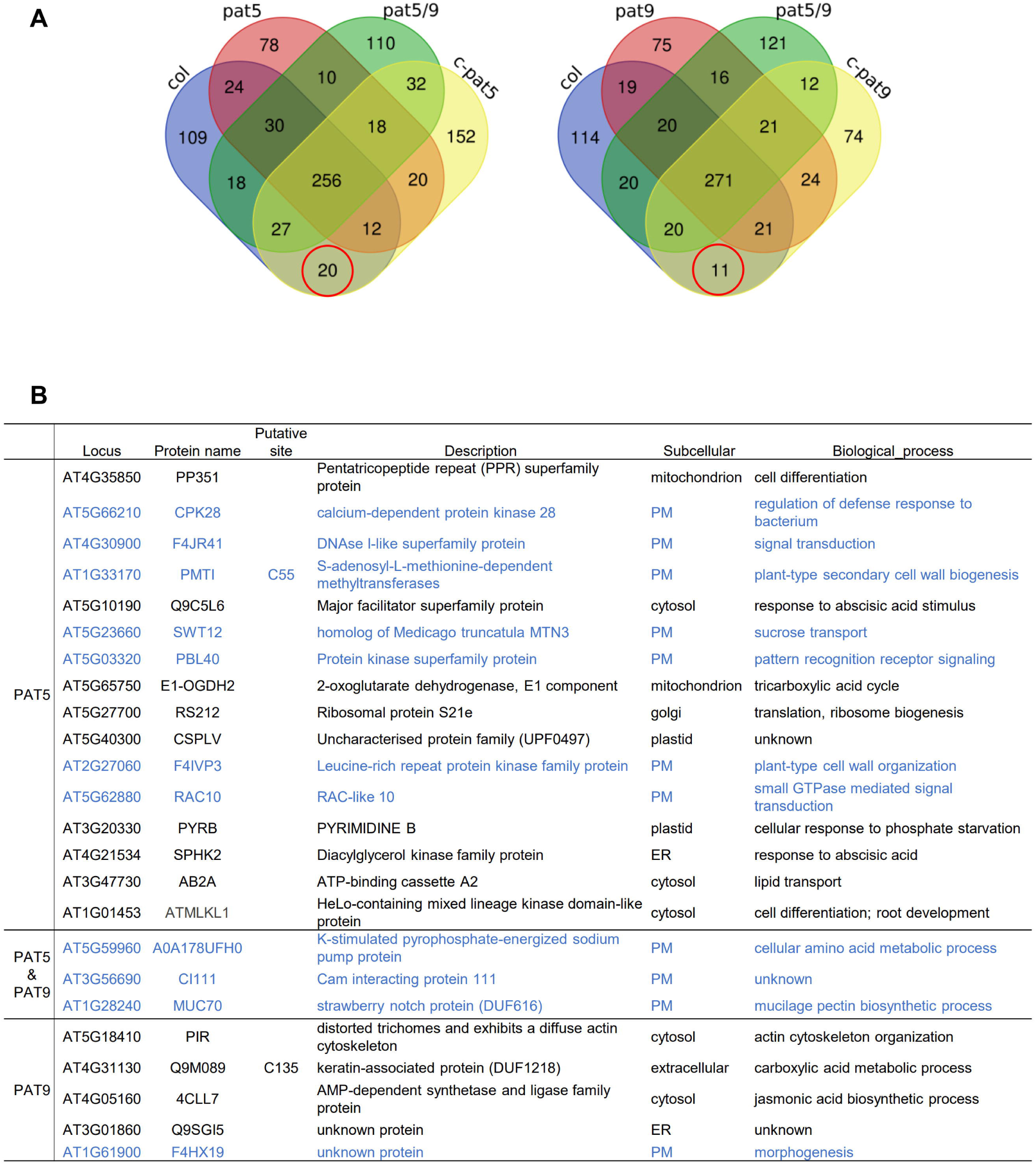
Identification of substrates of PAT5 and PAT9 using S-acyl-proteome. (A) Venn diagram for S-acylated proteins identified in each genotype (Col-0, *pat5*, *pat5*/*pat9*, c-*pat5*, *pat9*, c-*pat9*) using ABE-protein method. Candidate substrates for PAT5 and PAT9 are indicated with red circles. (B) Annotations for candidate substrates of PAT5 or PAT9. Locus duplicates from different Uniprot accessions were removed. Plasma membrane localized proteins were highlighted in blue color.

Meanwhile, 120 proteins (inferred from 139 peptides) and 114 proteins (inferred from 114 peptides) were identified to be potential substrates of PAT5 and PAT9, respectively, from the ABE-peptide S-acylproteomes (Figure S8, Table S6). Among them, 31 proteins were potential substrates for both PAT5 and PAT9. PM localized candidate substrates included three receptor-like kinases (CRK10, NILR2, SIRK1), disease response proteins (RIN4, RPP8), and vesicle-associated membrane proteins (VAMP722, CLAH2, DRP1A, SEC3A, SEC5A, AP2M, CLC2, DRP2B). These potential substrates suggested that in addition to immune responses, PAT5 and PAT9 may regulate multiple biological processes through S-acylating their substrates. The only functionally characterized substrate P2K1 of PAT5 and PAT9 (Chen et al. 2021) was not detected in this study. We attribute this to the low abundance of P2K1, as shown by independent LC-MS analysis (i.e., P2K1 peptides were not detected in the global proteome of the ePM samples). No candidate substrates from the ABE-peptide data overlapped with those from the ABE-protein data.

Although we were not able to detect P2K1 through proteomics in WT plants, an independent S-acylproteome using the ABE-protein method with tissue derived from a P2K1 overexpressing (35S::P2K1/Col) transgenic line showed significant enrichment of P2K1 in the +Hyd sample, indicating an S-acylated state of P2K1. To further characterize S-acylation sites of P2K1, an immunoprecipitation (IP)-MS based S-acylation detection experiment was conducted using stable transgenic NP::P2K1-HA/Col-0 plants. C407 site (Figure S9) was detected as a putative S-acylation site, which was previously identified to be S-acylated through mutagenesis and biochemical experiments (Chen et al. 2021). Thus, our proteomic data validated the S-acylation of P2K1 *in vivo*.

### Experimental verification of the S-acyl proteome

To further evaluate the quality of our proteome data, the ABE and immunoblotting assays were used to experimentally verify the S-acylation of the following eight candidate proteins: Five candidate proteins from the RLK or RLCK family (FERONIA, BIR2, CARK8, BSK3, and BSK5) and three from PAT5 or PAT9 candidate substrates (CPK28, PBL40, and RIN4). The selected candidate proteins were transiently expressed in tobacco to verify their S-acylation states. As shown in Figure 9, after ABE enrichment, all the tested proteins were enriched under the Hyd+ condition compared with the Hyd− condition, indicating that our S-acylation data are of high quality.

**Figure 9.**
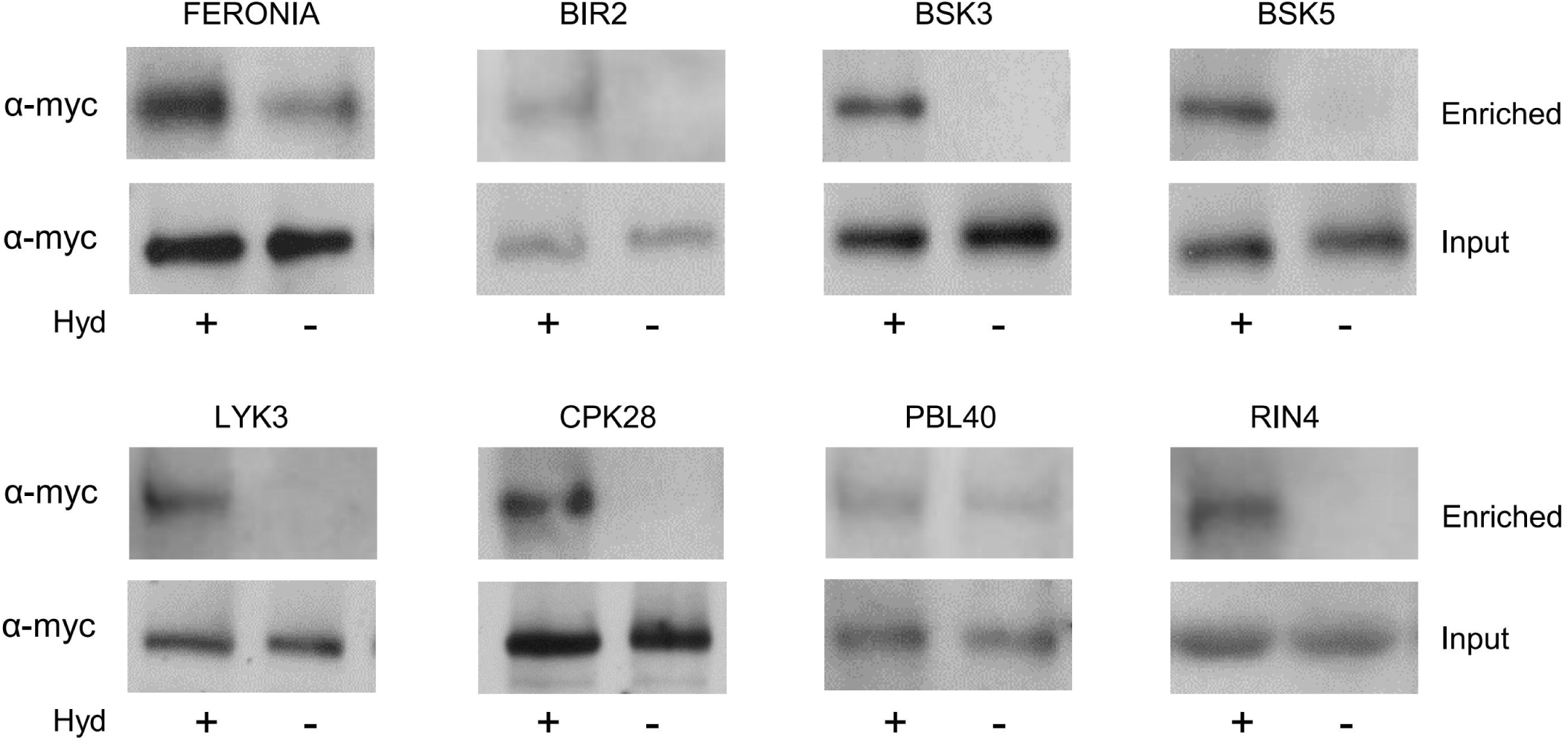
Validation of S-acylated proteins using ABE and immunoblotting method. ABE assay was performed with extracts containing myc-fusion proteins transiently expressed in tobacco using 35S promoter and immunoblots were probed with anti-myc antibody. Hyd indicates presence (+) or absence (−) of hydroxylamine required for S-acylation enrichment. The enriched sample was compared with the input which was used as loading control.

## Discussion

In this study, we employed two S-acylation enrichment techniques: the ABE-protein and ABE-peptide methods, to capture the *Arabidopsis* S-acylproteomes, with a focus on membrane proteins. The two methods produced distinct yet complementary S-acylproteomes, likely due to intrinsic detection biases associated with each method. Notable differences were observed between these two methods in terms of the identification of S-acylation sites, as well as the total number and diversity of S-acylated proteins detected. The ABE-protein method tended to enrich and reveal S-acylated proteins from the plasma membrane, while the ABE peptide method identified more S-acylated proteins from the cytosol. We speculate that proteins with intensive S-acylations, e.g., double or triple consecutive S-acylations in many PM proteins, may be better captured by the ABE-protein method due to their stronger binding affinity to the capture resin. S-acylated proteins with high protein abundance, predominantly cytosolic proteins, are also likely to be captured by the resin. Consequently, these proteins are more readily detected by the ABE-peptide method. Notably, the ABE-peptide method revealed many more potential S-acylated proteins than the ABE-protein method. Additionally, the ABE peptide method was able to reveal the S-acylation sites directly. Therefore, by combining these two approaches, a better picture of the protein S-acylation at the proteome level can be achieved in an organism. Of course, any method may be prone to issues of partial coverage and false identification. For example, some S-acylated proteins may be too low in abundance to be confidently revealed by the current methods. Additionally, the S-acylation of some other proteins may be regulated by different growth stages, tissues/organs, and environmental stimuli. Therefore, more detailed work is needed to reveal a complete S-acylproteome for *Arabidopsis*.

By integrating the data obtained using these two methods, our study successfully revealed a comprehensive and well-represented S-acylproteome for *Arabidopsis*, especially its membrane proteins. Together with the PAT mutants and complementation transgenic lines, over 2000 S-acylated proteins and 2500 S-acylation sites were identified. It is estimated that over 10% of the proteome in mammalian systems may be S-acylated (Edmonds et al. 2017). Applying this estimation to *Arabidopsis*, one could anticipate over 2500 S-acylated proteins in this organism. Therefore, our current study may have revealed a significant portion, if not the entirety, of the total S-acylated proteins in *Arabidopsis*.

Our data showed significant overlap with two previous S-acylproteomics studies conducted in *Arabidopsis* using different methods and tissues (Hemsley et al. 2013, Kumar et al. 2022), underscoring the high quality of our results. In the S-acylproteomics study by Hemsley *et al*. (2013), 581 putative S-acylated proteins were identified from *Arabidopsis* root suspension cells by protein-level enrichment method coupled to the isobaric tagging for relative and absolute quantitation (iTRAQ)-based LC-MS. Less strict criteria (fold change of +Hyd/-Hyd > 1, without p value from t-testing as a criterion) were used in this study. Due to the limitation of the method, no S-acylation sites were found. In the study by Kumar *et al*. (2022), many S-acylated proteins and sites were identified from multiple tissues (i.e., stem, hypocotyl, silique, meristem, seedling, leaf) by an acyl-RAC based peptide-level enrichment approach. The total number of identified S-acylated proteins in our ABE-peptide dataset is similar to that discovered in seedlings, but is lower than the number reported from all six tissues in the above acyl-RAC study, suggesting tissue-specific S-acylation for some proteins. Despite this, there was a significant overlap between our identified S-acylated proteins and those reported in the aforementioned studies. The many proteins not overlapping with these previous studies may be novel S-acylated proteins, thus expanding the *Arabidopsis* S-acylproteome. Furthermore, our emphasis on membrane-enriched fractions led to the identification of a substantial number of S-acylated proteins that are likely involved in signal transduction and environmental sensing.

Prior to our study, no clear patterns of protein S-acylation had been identified. We discovered that the surrounding amino acids, the relative position of the cysteine residue within a protein, and the protein’s subcellular localization significantly influence the S-acylation status of a particular cysteine residue. While motif analysis did not reveal clear conserved sequences around the S-acylation sites, certain surrounding amino acids exhibited a significant overrepresentation or underrepresentation, especially in certain groups of proteins. Although S-acylation sites were generally found throughout the entire length of a protein, they were unevenly distributed within a protein, likely due to the asymmetrical distribution of cysteine residues and their surrounding amino acids. Interestingly, proteins from different organelles showed different distribution patterns of S-acylated cysteines. For example, S-acylated proteins predicted to localize to mitochondria showed intensive S-acylation at two ends, while proteins predicted to be in the vacuole showed an opposite pattern. These findings suggest that the S-acylation status of a cysteine residue is controlled by multiple factors and more work is needed to understand the impact of these factors on S-acylation and the corresponding biological functions.

Prior to our study, only a few substrate proteins had been identified for a few PATs, e.g., ROP2 for PAT4 (Wan et al. 2017). By utilizing S-acylproteomics data from both wild-type plants and PAT5 and PAT9 mutants, we identified many potential substrates for these two key plant immunity-related protein S-acyltransferases. These identified substrates are involved not only in defense responses (CDPKs, PBL40, RIN4, RPP8), but also in cellular signal transduction (RAC10, BON2), and cell wall biogenesis (PMTI, F4IVP3), suggesting that PAT5 and PAT9 may regulate many other biological processes in addition to immunity. Unfortunately, the only functionally characterized substrate P2K1 of PAT5 and PAT9 (Chen et al. 2021) was not detected in our study using WT *Arabidopsis* Col-0, likely due to its low abundance. However, we were able to confirm *in vivo* the previously identified C407 S-acylation site through analysis of the P2K1 over-expression plants. Therefore, our study not only revealed potential substrates for the two important defense-related PATs, but also indicated that a single PAT may S-acylate multiple proteins and a single protein may be S-acylated by multiple PATs. This indicates that the regulation of S-acylation in plants is likely complex, and additional studies are needed to dissect PAT-substrate relationships, especially under different internal and external stimuli.

In summary, the combined use of our optimized ABE-protein and ABE-peptide methods proved effective in identifying a broad range of S-acylated proteins. Through the application of these two complementary methods, our study not only significantly expanded the *Arabidopsis* S-acylproteome, but also revealed potential substrates for the two key defense-related PATs. These findings provide valuable insights into the understanding of protein S-acylation in plants and serve as an invaluable resource for the research community to further study the functions of S-acylated proteins.

## Materials and Methods

### Plant Materials and Growth Conditions

The *Arabidopsis thaliana* mutant lines *pat5* (GABI-322D08; At3g48760), *pat9* (SALK-003020C; At5g50020), and the double mutant *pat5*/*pat9* are in the Columbia (Col-0) background. The complemented transgenic lines of *pat5* and *pat9* mutants, NP::PAT5/*pat5* (c-*pat5*) and NP::PAT9/*pat9* (c*-pat9*), and the P2K1-HA fusion transgenic line NP::P2K1-HA/Col-0 were previously generated (Chen et al. 2021). Sterilized *Arabidopsis* seeds were germinated and grown on 1/2 MS medium containing 1% sucrose at 23L, 60%-70% humidity in a growth chamber with a 16-hour light/8-hour dark cycle. After 13 days, seedlings were transferred to deep petri dishes with 1/2 MS liquid medium and placed in the growth chamber overnight. After removing excess water with paper tissue, 14-day-old seedlings were immediately frozen in liquid nitrogen and stored at −80L until further use. Four independent biological replicates for each genotype were prepared for both protein-level and peptide-level enrichment methods.

### Isolation and enrichment of plasma membrane proteins

Plasma membrane proteins from the 14-day-old seedlings of six *Arabidopsis* genotypes (Col-0, *pat5*, *pat9*, *pat5*/*pat9*, c-*pat5* and c-*pat9*) were isolated and enriched as previously described (Zhou et al. 2020). Briefly, Seedlings were ground into a fine powder in liquid nitrogen and then reconstituted in ice-cold Homogenization buffer (H buffer, 50Mm HEPES pH7.5, 250mM sucrose, 5% (v/v) glycerol, 10mM EDTA pH 8.0, 0.5% (w/v) polyvinyl pyrrolidone, proteinase and phosphatase inhibitors) in 50-ml tubes. The samples were centrifugated at 10,000 g at 4L for 10 min, and the supernatants were filtered through Miracloth. The filtered supernatants were further centrifugated at 121,000 g at 4L for 4 hours to obtain crude membrane pellets. The crude membrane pellets were resuspended in H buffer plus 0.2% Brij-58 (1 μL buffer per 5 μg crude membrane proteins) in 1.5-ml safe-lock tubes and incubated on ice for 30 min. The resultant samples were centrifugated for 30 min at 100,000 g at 4L. The pellets were further treated with Brij-58 and subjected to ultracentrifugation as described above to obtain enriched plasma membrane (ePM) fractions. Protein concentrations were determined using the BCA assay.

### S-acylation Enrichment by Acyl-biotinyl Exchange (ABE) at the Protein-level and Peptide-level

About two mg enriched plasma membrane (ePM) proteins of aforementioned six genotypes underwent the S-acylation enrichment according to the ABE-protein and the ABE-peptide methods described by Zhou *et al*. (Zhou et al. 2020). The ePM proteins were precipitated using the methanol/chloroform precipitation method and then re-solubilized in 2% SDS buffer (2SB buffer, 50mM Tris-HCl pH7.4, 2% (w/v) SDS, 5mM EDTA). TCEP (Sigma-Aldrich, cat. no. C4706) was added to a final concentration of 50 mM and incubated at room temperature (RT) for 1 hour with rotation to break the disulfide bonds. After the reduction reaction, excess TCEP was removed by methanol/chloroform precipitation. Protein pellets were resolubilized in 2SB buffer as described above. To alkylate the free thiol, NEM (Sigma-Aldrich, cat. no. 04259) was added to a concentration of 50 mM and incubated in the dark at 55°C with shaking for 1 hour. Excess NEM was removed by methanol/chloroform precipitation. Protein pellets were resolubilized in 2SB buffer. DTDP (Sigma-Aldrich, cat. no. D5767) was added to a final concentration of 25 mM to further block free thiol. Excess NEM and DTDP were removed by two sequential methanol/chloroform precipitations. Proteins were resolubilized in 2SB buffer and divided equally into two samples: the experimental and control samples. To the experimental sample (+Hyd), hydroxylamine (Hyd, Sigma-Aldrich, cat. no. 159417) and HPDP-biotin (Thermo Fisher, cat. no. 21341) were added to a final concentration of 0.5 M and 0.25 mM, respectively. To the control sample (-Hyd), 50mM Tris-HCl and HPDP-biotin were added. Both samples were incubated at RT for 1 hour with rotation. Excess HPDP-biotin was removed by methanol/chloroform precipitation.

For protein-level enrichment, protein pellets were resolubilized in a small volume of 2SB buffer. After complete solubilization, 19 volumes of Dilution buffer (D buffer, 50 mM Tris-HCl pH7.4, 150 mM NaCl, 5 mM EDTA, 0.2% (v/v) Triton X-100) were added to decrease the concentration of SDS to 0.1%. After vortexing and then centrifugation at 16,000 g for 5 min, the supernatant was transferred to pre-equilibrated high-capacity streptavidin agarose resin (Pierce, cat. no. 20357). The mixture was incubated with rotation at RT for 1 hour, and then transferred to a spin column, followed by centrifugation for 1 min at 400 g. The resin was thoroughly washed six times with the following Equilibration buffer (E buffer) by inverting, followed by centrifugation for 1 min at 400 g. Equilibration buffer: E buffer, 50 mM Tris-HCl pH7.4, 150 mM NaCl, 5 mM EDTA, 0.2% (v/v) Triton X-100, 0.1% (w/v) SDS. To elute the enriched proteins, 50 mM TCEP in E buffer was added to the column with the bottom cap on and incubated at 37°C with shaking for 20 min. The columns were inserted into 1.5 ml tubes and centrifuged at 400 g for 2 min to collect the enriched proteins. Unlabeled cysteines (putative S-acylation sites) were then alkylated by adding 500 mM iodoacetamide (IAM, Pierce, cat. no. A39271) to a final concentration of 10 mM and incubated in the dark for 45 min at room temperature. Alkylated proteins were then digested with the S-Trap^TM^ micro spin columns (PROTIFI, cat. no. C02-micro-10) following the protocols provided by the manufacturer (https://protifi.com/pages/protocols). In our study, trypsin was used at a 1:10 (w/w) trypsin-to-protein ratio and incubated at 47°C for 1 hour to digest proteins. Peptides were eluted, pooled and dried as described (Zhou et al., 2000), and resuspended in 0.1% formic acid (FA) in 3% acetonitrile (ACN) in water at 0.1 μg/μL for MS analysis.

For peptide-level enrichment, after labeling the S-acylation sites by the ABE process, biotinylated proteins were resolubilized in 50 μL of 8 M urea/100 mM NH_4_HCO_3_, pH 8 and then diluted with 100 mM NH_4_HCO_3_ to reduce the urea concentration to < 1 M. Un-solubilized protein aggregates were removed by centrifugation for 5 min at 16,000 g. Proteins were then digested with trypsin at a 1:50 trypsin-to-protein ratio overnight at 37°C. 2 × E buffer were added to achieve a final buffer with 50 mM Tris-HCl (pH 7.4), 150 mM NaCl, 5 mM EDTA, 0.1% SDS, 0.2% Triton X-100, 0.4 M urea, and 50 mM NH_4_HCO_3_. To capture the biotinylated peptides, 100 μL equilibrated streptavidin agarose resin was added to each sample and incubated at RT for 60 min with rotation. The resins were washed with E buffer for three times and washing buffer (2 M urea in 100 mM Tris pH7.4) for another three times. Peptides were eluted with the elution buffer (50 mM TCEP in 100 mM Tris-HCl, pH7.4) by incubating at 37°C with rotation for 20 min and centrifugation for 2 min at 400 × g. A final concentration of 10 mM IAM was added and incubated in the dark at RT for 45 min to alkylate the peptides. 2-disks CDS Empore C18 (CDS Analytical) StageTips were conditioned and equilibrated with 100 μL acetonitrile (ACN), 100 μL 50% ACN/0.1% FA, and 200 uL 0.1% FA. After equilibration, peptides were loaded onto conditioned StageTip, washed with 200 μL 0.1% FA and eluted with 50 μL 50% ACN/0.1% FA. Desalted peptides were transferred to HPLC vials and dried via vacuum centrifugation. Peptides were then resuspended in 0.1% FA in 3% ACN in water at 0.1 μg/μL for MS analysis.

### Immunoprecipitation (IP)-MS based Identification of S-acylation Sites

To identify the S-acylation sites of the low abundant protein P2K1, the only reported substrate of PAT5 and PAT9, an IP-MS based S-acylation detection experiment was conducted using stable transgenic NP::P2K1-HA/Col-0 plants. Total proteins were extracted from 14-day-old seedlings with 10 mL Radio Immunoprecipitation Assay lysis buffer (RIPA buffer, 50 mM Tris-HCl (pH7.4), 150 mM NaCl, 1% (v/v) Nonidet P-40 (NP-40), 1 mM EDTA, 0.5% (w/v) sodium deoxycholate, 0.1% (w/v) SDS, protease inhibitor cocktail) by incubating at 4 °C > 1 hour with rotation. After centrifugation at 12,000 g for 5 min at 4 °C, the supernatant was transferred and centrifugated again. The supernatant was transferred to a fresh tube and placed on ice for immunoprecipitation. Two hundred microliters of anti-HA-agarose suspension was added to the supernatant and incubated overnight with rotation at 4 °C. After a short spin, the supernatant was discarded, and the rest of the anti-HA-agarose suspension was transferred to a spin column. The resin was washed for 3 times with RIPA buffer and pelleted by spinning at 300 g for 1 min. Flow-through was discarded and the column was plugged. 10 mM Tris (2-carboxyethyl) phosphine hydrochloride (TCEP) in the Washing buffer was added to the column and incubated for 1 hour with gentle rotation at RT. Flow-through was discarded and the resin was washed with the Washing Buffer.

To block the free Cys, freshly prepared 50 mM N-ethylmaleimide (NEM) was added to the column and incubated in the dark for 1 hour with gentle rotation at RT. The resin was then washed with the Washing Buffer for 3 times and then divided equally into two samples. The experimental sample was incubated in 1 M hydroxylamine (Hyd) for 1 hour to cleave the thioester bonds of S-acylation sites, and the control sample was incubated with 50 mM Tris-HCl (pH7.4). After washing the resin with the Washing Buffer, 10 mM IAM was added and incubated for 1 hour at RT. The proteins were finally eluted by adding 100 µL Elution buffer (50 mM Tris-HCl (pH7.4), 2% (w/v) SDS, 10 mM TCEP) and heating at 75 °C for 20 min. The eluted proteins were digested with trypsin. The generated peptides were desalted with C18 tips and then resuspended in 0.1% FA in 3% ACN in water at 0.1 μg/μL for MS analysis.

### LC-MS/MS Analysis

Digested samples were analyzed using Waters nanoACQUITY UPLC System coupled with Thermo Orbitrap Fusion Lumos mass spectrometer. LC system was configured with Mobile phase A (MPA, 0.1% (v/v) formic acid and 3% ACN in water) and Mobile phase B (MPB, 0.1% (v/v) formic acid and 10% water in ACN) at 0.2 μL/min. Samples were directly loaded to the in-house packed analytical column (Dr. Maisch ReproSil-Pur 120 C18 1.9 μM particle, column ID 75 μM, OD 360 μM, length ∼20 cm, heated at 50 °C) for 30 min at 2% MPB. Then the gradient started from 6%-30% over 84 min, followed by ramp to 90% MPB over 10 min and subsequent equilibration. The Orbitrap instrument was operated in data-dependent acquisition mode with the following settings. Ion source: electrospray voltage 2.2 kV, s-lens RF 30%, capillary temperature 250°C, no source gas; MS1: 60,000 resolution, automatic gain control (AGC) target 4e5, maximum injection time 50 ms, scan range 350-1800 m/z, profile spectra; MS2: 50,000 resolution, AGC target 1e5, maximum injection time 105 ms, isolation window 0.7 m/z, normalized collision energy 30%, centroid spectra; data-dependent acquisition: cycle time 2 s, intensity threshold 2.5e4, include charge state 2 - 6, dynamic exclusion of 45 s. 5 μL peptide solution (0.1 μg/μL) was injected for each sample and the MS data was collected.

### Database Searching

MS/MS spectra were searched against the *Arabidopsis thaliana* proteome sequences downloaded from UniProt database (https://www.uniprot.org/) using the PEAKS Studio 10.0. Various modifications, such as carbamidomethyl on Cys, N-ethylmaleimide on Cys, dithiodipyridine on Cys, methionine oxidation, and protein N-terminal acetylation, were included in the peptide search. A maximum of 3 modifications were allowed per peptide. Among them, carbamidomethyl on Cys is the potential S-acylation site. The precursor mass tolerance of 10 ppm and the fragment mass tolerance of 0.02 Da was used. The false discovery rate (FDR) for peptide, protein, and PTM identifications was set to 0.01. The peptide-level data were also processed with MaxQuant (v2.0.1.0) with same modification settings and default label-free quantitation parameters.

### Bioinformatic Analysis

Initial data processing was performed using the R programming language. The DEP package (Zhang et al. 2018) from Bioconductor was used to conduct data preparation, as well as identification of enriched proteins/peptides. Intensity values of proteins/peptides were used to quantify abundance and enrichment of proteins/peptides. Data was filtered so that proteins/peptides were identified in at least 2 out of 4 replicates under at least one condition. Raw intensities were log2 transformed, and normalized by variance stabilizing transformation. Missing values were imputed using random draws from a Gaussian distribution centered around a minimal value. A PCA analysis for all the samples was performed using protein/peptide intensities to evaluate the reproducibility between replicates. Based on the PCA analysis, two samples (*pat5* -Hyd and *pat5*/*pat9* -Hyd) from the protein-level enrichment set and the fourth replicate in the peptide-level enrichment set (batch effect) were considered to be outliers and removed from further analysis. Fold change (FC) and FDR value were calculated between the +Hyd/−Hyd treated samples to quantify enriched proteins/peptides. Proteins identified in protein-level enrichment or peptides identified in peptide-level enrichment method enriched with at least a 2-fold change and an FDR within 0.05 were defined as being enriched and considered to be putative S-acylated proteins/peptides.

Analysis of GO term enrichment in the set of S-acylated proteins compared to the *Arabidopsis* genome was performed using the PANTHER overrepresentation test (Mi et al. 2005, http://geneontology.org/). Protein subcellular localizations were analyzed using SUBA4 (Hooper et al. 2017, https://suba.live/). The enrichment p-value of protein subcellular localization was calculated based on the hyper-geometric distribution test. An S-acylation site motif analysis was performed using the pLogo generator (O’Shea et al. 2013, https://plogo.uconn.edu/). 15 amino acid peptide sequences containing the central cysteine and 7 flanking amino acids on each side were loaded as the foreground. The *Arabidopsis* proteome was selected as the background. The significance threshold was set to p < 0.05. STRING (Szklarczyk et al. 2021, https://cn.string-db.org/) was used for protein-protein interaction pathway analysis and only interactions with high confidence (minimum interaction score > 0.7) were shown.

### Validation of candidate S-acylated proteins using ABE assay and immunoblotting

To validate the S-acylation of candidate proteins, myc-fusion protein constructs were made and transiently expressed in *Nicotiana benthamiana* leaves, followed by the ABE-protein based S-acylation enrichment and immunoblotting assay. Briefly, the full-length coding sequences of the selected candidate genes were amplified from wild-type plants. PCR products were cloned into the entry vector pDONR207 using BP cloning and then cloned into the destination vector pGWB617 using LR cloning. The confirmed gene constructs were introduced into 4-week-old *Nicotiana benthamiana* leaves using *Agrobacterium* infiltration. Twenty-four hours after infiltration, the injected tobacco leaves were collected. The total protein was isolated and the ABE-protein assay was conducted as described above. Instead of trypsin digestion, proteins were eluted with the TCEP buffer and separated on 10% SDS-PAGE gels, blotted onto PVDF membranes, and probed with anti-myc antibody (Abmart, Cat#M20019, diluted at 1:2000).

## Supplementary Data

**Supplemental Figures**

**Supplemental Tables**

## Data availability

All MS data are available at MassIVE (massive.ucsd.edu) with accession MSV000088798 (for the reviewer, data can be accessed via FTP using Server: massive.ucsd.edu; User: MSV000088798; Password: Plant4000).

## Acknowledgements

Not applicable.

## Author Contributions

G. S. and L. Z. conceptualization; L. Z., and M. Z. methodology; L. Z., M. Z., M. A. G., Y. M., and Y. Z. investigation; L. S. and M. Z. software; L. Z., L. S., and M. Z. formal analysis; M. Z. data curation; G. S. and L. P.T. funding acquisition; G. S., L. P.T., and D. X. supervision; L. Z. writing - original Draft; G. S., J. W., M. Z., L. P.T., and M. A. G. writing - review & editing.

