## Supplemental Figures for "Global analysis of membrane protein S-acylation in the model plant *Arabidopsis thaliana*"

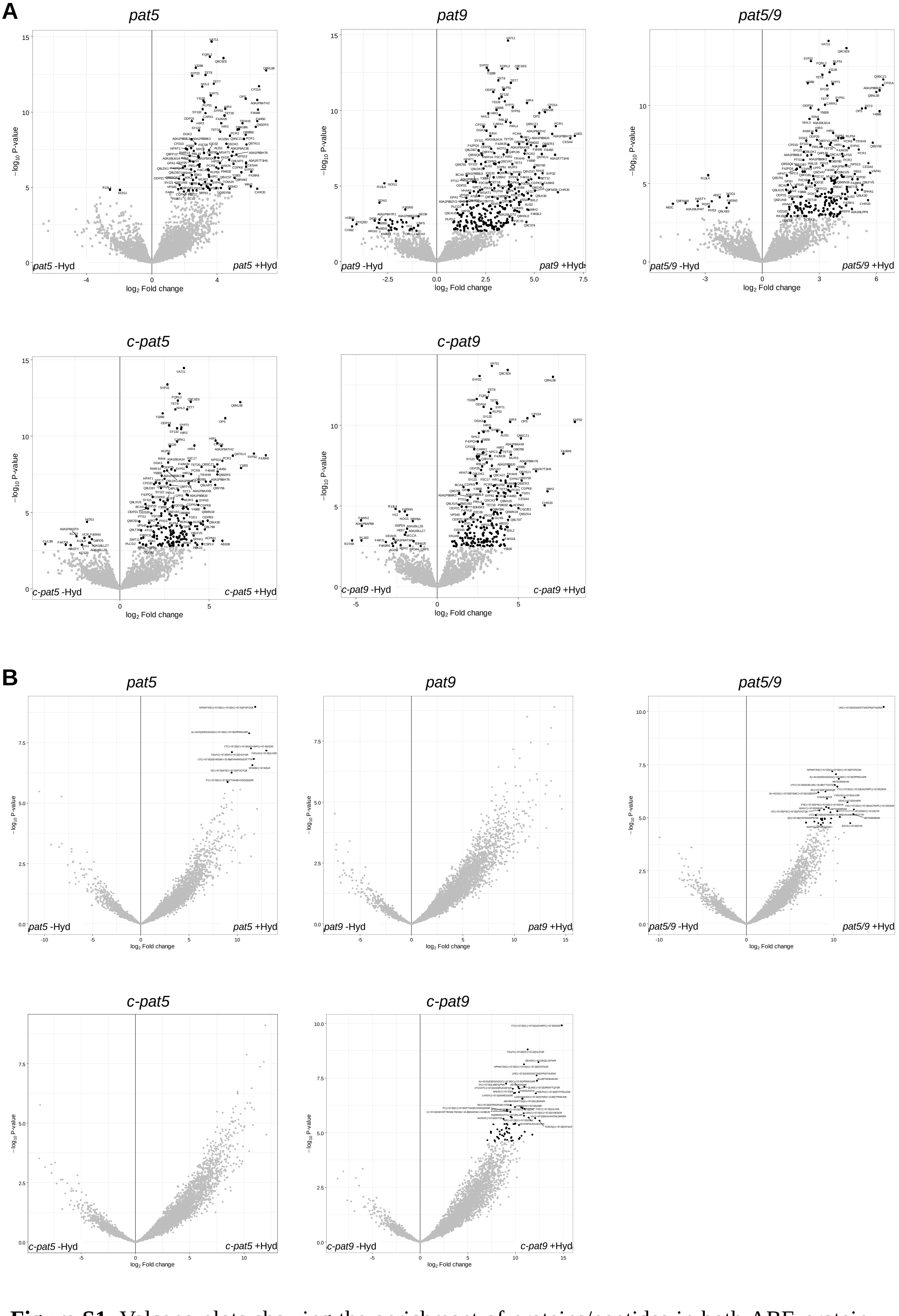


**Figure S1.** Volcano plots showing the enrichment of proteins/peptides in both ABE-protein (A) and ABE-peptide (B) S-acylproteomes from *pat5*, *pat9*, *pat5/pat9*, c-*pat5* and c-*pat9*.

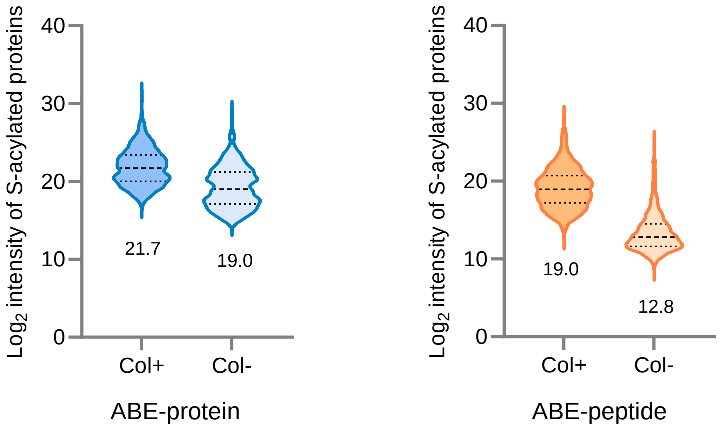


**Figure S2.** Violin plot of log2 transformed intensities of the enriched proteins identified in the ABE-protein method (left) and the enriched peptides identified in the ABE-peptide method (right) from Col-0.

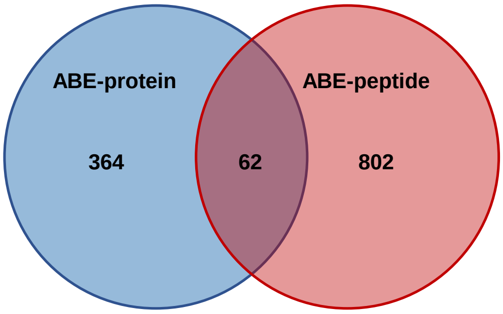


**Figure S3.** Overlap of S-acylated proteins identified in each method from Col-0.

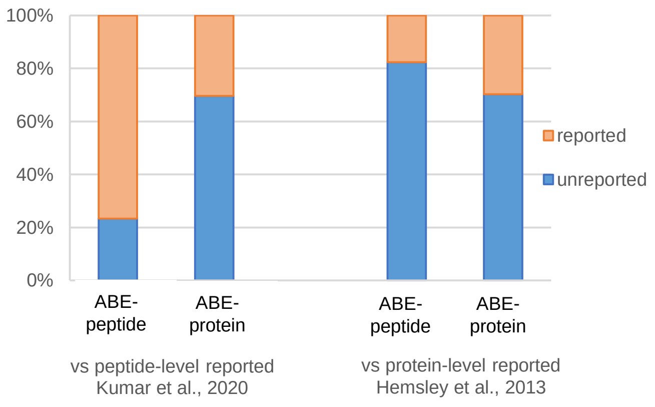


**Figure S4.** Proportion of previously reported and newly discovered S-acylated proteins identified in our study.

| Location | database annotated | ePM total | ABE-protein | | | ABE-peptide | | |
|---|---|---|---|---|---|---|---|---|
| | | | all detected | enriched | none-enriched | all detected | enriched | none-enriched |
| extracellular | 10.9% | 6.2% | 5.2% | 4.2% | 6.0% | 7.2% | 5.8% | 7.6% |
| PM | 11.7% | 14.2% | 14.3% | 41.1% | 13.6% | 12.7% | 13.2% | 13.8% |
| cytosol | 19.7% | 15.8% | 19.6% | 11.0% | 16.0% | 19.5% | 22.0% | 18.2% |
| nucleus | 27.9% | 7.3% | 12.0% | 8.2% | 25.4% | 16.0% | 7.5% | 24.5% |
| ER | 2.7% | 8.8% | 6.5% | 3.1% | 4.8% | 4.2% | 4.8% | 2.2% |
| golgi | 2.2% | 8.9% | 7.8% | 9.6% | 4.7% | 6.6% | 7.5% | 3.7% |
| plastid | 9.9% | 17.1% | 16.1% | 7.3% | 13.6% | 16.7% | 19.1% | 14.7% |
| mitochondrion | 8.3% | 10.3% | 8.3% | 4.9% | 8.1% | 7.3% | 8.5% | 7.3% |
| vacuole | 1.9% | 3.6% | 2.7% | 5.6% | 2.3% | 2.4% | 2.8% | 1.4% |
| peroxisome | 1.2% | 2.2% | 2.1% | 0.7% | 1.7% | 2.0% | 2.0% | 1.7% |
| others | 3.7% | 5.7% | 5.4% | 4.2% | 3.8% | 5.3% | 6.9% | 4.7% |

**Figure S5.** Percentage of protein subcellular localizations of each protein category in the S-acyl-proteomes. Compared to the ePM global proteome, the subcellular enrichment of the protein categories in the S-acyl-proteomes was tested using the hyper-geometric distribution. Significantly changed localizations ($p < 0.001$) are indicated with red (enriched) and blue (decreased) shades.

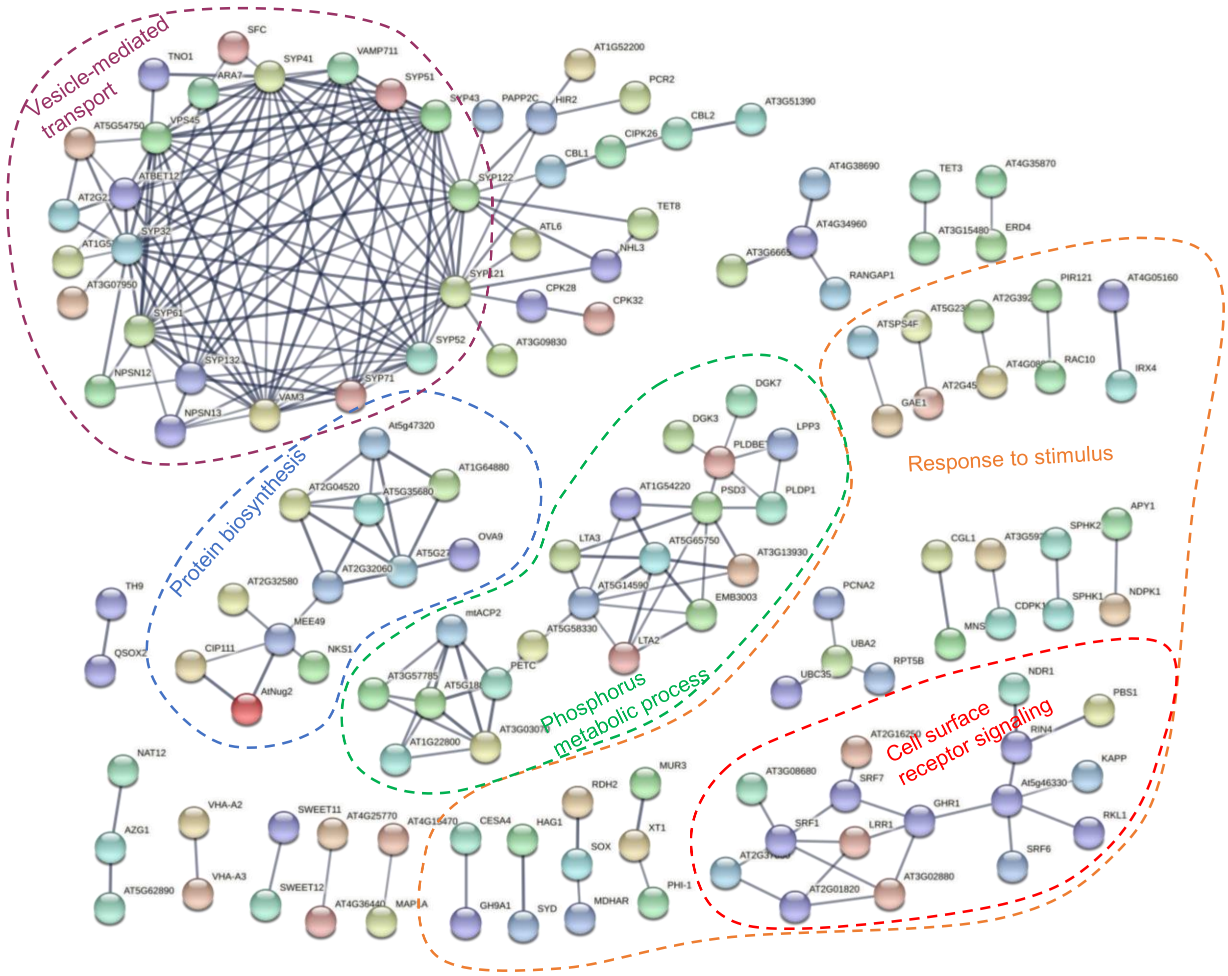


**Figure S6.** Protein-protein interaction (PPI) analysis using STRING. S-acylated proteins identified by the ABE-protein method were used for analysis and only PPIs with high confidence (interaction score > 0.7) are shown. For simplicity, the functions of the scattered PPIs were not specified in the figure.

| RLK family | Locus | Protein. name | Identified approach 1. ABE-protein 2. ABE-peptide | Protein length | S-acylation sites | Col-0 | *pat5* | *pat9* | *pat5/9* | *c-pat5* | *c-pat9* |
|---|---|---|---|---|---|---|---|---|---|---|---|
| CrRLK1L-1 | AT3G51550 | FERONIA | 2 | 895 | C521, C639 | 7.1 | 2.1 | 4.2 | 6.3 | 8.3 | 4.2 |
| CrRLK1L-1 | AT5G38990 | MDS1 | 2 | 880 | C281, C510 | 4 | -0 | 0.7 | 4.7 | 1.1 | 3.3 |
| CrRLK1L-1 | AT5G54380 | THE1 | 1 | | | 2.5 | 2.4 | 2.2 | 4.2 | 2.4 | 3.4 |
| DLSV | AT4G23250 | A0A1P8B8K3 | 1 | | | 2.4 | 2.4 | 2 | 2.2 | 2.6 | 2.1 |
| DLSV | AT1G16670 | CRPK1 | 1, 2 | 390 | C147, C296 | 1.8 | 2 | 1.4 | 2.2 | 2.3 | 1.9 |
| LRK10L-2 | AT1G66930 | Q9FZI4 | 1 | | | 3.3 | 4.5 | 3.7 | 3.1 | 2.8 | 4.1 |
| LRR-I-1 | AT2G37050 | C0LGM1 | 1 | | | 2.2 | 2 | 2.4 | 2.3 | 1.6 | 2.7 |
| LRR-I-1 | AT1G51800 | IOS1 | 2 | 894 | C836, C857 | 3.2 | 1.8 | 1.5 | 1.8 | 1.8 | -0 |
| LRR-II | AT5G65240 | F4KGL1 | 1 | | | 2.4 | 2.4 | 4.2 | 2.4 | 3.6 | 3.1 |
| LRR-II | AT1G71830 | SERK1 | 2 | 625 | C366 | 2.9 | 4.9 | 1.2 | 6.2 | -1 | 3.4 |
| LRR-II | AT5G10290 | Y5129 | 1 | | | 1.6 | 1.6 | 2.1 | 2.1 | 2.2 | 2.2 |
| LRR-III | AT2G27060 | F4IVP3 | 1 | | | 2.6 | 1.3 | 2 | 1.2 | 2.4 | 0.6 |
| LRR-III | AT4G20940 | GHR1 | 1 | | | 2.3 | 2 | 1.6 | 2.4 | 0.9 | 1.3 |
| LRR-III | AT3G56100 | IMK3 | 1 | | | 2.4 | 0.9 | 2.3 | 1.4 | 0.2 | -0 |
| LRR-III | AT3G02880 | KIN7 | 1, 2 | 627 | C217, C594 | 2.2 | 2.5 | 2.6 | 2.4 | 2.4 | 2.4 |
| LRR-III | AT5G16590 | LRR1 | 1, 2 | 625 | C488, C592 | 5.5 | 3.9 | 3.4 | 4.8 | 5.8 | 6.1 |
| LRR-III | AT3G17840 | RLK90 | 2 | 647 | C222, C229 | 4.3 | 5.2 | 4.1 | 5.5 | 5.8 | 1.9 |
| LRR-III | AT5G10020 | SIRK1 | 2 | 1048 | C204, C342 | 5.1 | 2.1 | 3 | 1.9 | 2.2 | 8.1 |
| LRR-III | AT1G48480 | Y1848 | 1 | | | 2.4 | 3.4 | 3 | 3 | 3.9 | 3.5 |
| LRR-III | AT3G08680 | Y3868 | 1 | | | 2.4 | 2.7 | 2.6 | 2.1 | 2.5 | 2.7 |
| LRR-IX | AT3G23750 | BARK1 | 2 | 928 | C356, C610, C836 | 1.9 | -1 | -0 | -1 | -2 | 0.2 |
| LRR-IX | AT2G01820 | TMK3 | 1 | | | 3.9 | 2.2 | 2.3 | 3 | 2.3 | 2.1 |
| LRR-V | AT2G20850 | SRF1 | 1 | | | 3.2 | 2 | -0 | 1.1 | 1.7 | -1 |
| LRR-V | AT1G53730 | SRF6 | 1 | | | 2.5 | 1.3 | 2.3 | 1.5 | 1.5 | 1.6 |
| LRR-V | AT3G14350 | SRF7 | 1 | | | 2.1 | 0.1 | 0.8 | -1 | 1 | 1.2 |
| LRR-VIII-1 | AT5G49760 | HPCA1 | 2 | 953 | C627, C883 | 2.3 | -1 | 1.8 | 3.1 | 3.7 | 3.1 |
| LRR-VIII-2 | AT1G16670 | CRPK1 | 1, 2 | 390 | C147, C296 | 3.3 | 3.2 | 5.7 | 5.1 | 7.4 | 4.2 |
| LRR-VIII-2 | AT3G14840 | LIK1 | 2 | 1020 | C776 | 5 | 6.6 | -1 | 4.5 | -1 | 1.3 |
| LRR-Xa | AT3G28450 | BIR2 | 2 | 605 | C101 | 7.5 | 3.7 | 4.7 | 5.2 | 6.3 | 5.8 |
| LRR-Xa | AT1G27190 | BIR3 | 2 | 601 | C30, C144, C334 | 5 | 3.7 | 3 | 1.1 | 1.3 | 2.3 |
| LRR-Xb-1 | AT4G39400 | BRI1 | 2 | 1196 | C268, C1137 | 4.8 | 4.2 | 4.9 | 0.6 | 3.5 | 2 |
| LRR-XI-1 | AT4G08850 | MIK2 | 1 | | | 1.4 | 1.6 | 1.5 | 1.7 | 2 | 1.6 |
| LRR-XII-1 | AT5G46330 | FLS2 | 1 | | | 3.5 | 3.3 | 3.6 | 2.7 | 2.7 | 3.3 |
| LRR-XIV | AT2G16250 | Y2165 | 1 | | | 2.5 | 1.1 | 2.3 | 1.8 | 0.7 | 2.5 |
| LysM | AT1G51940 | LYK3 | 1, 2 | 651 | C287 | 11 | 7.3 | 5 | 8.2 | 8.2 | 9.3 |
| LysM | AT2G23770 | LYK4 | 1 | | | 1.5 | 1.6 | 0.9 | 1.7 | 2.6 | 1.2 |
| PERK-1 | AT3G24550 | PERK1 | 1 | | | 1.7 | 1.5 | 2.2 | 2.4 | 3.7 | 2.2 |
| RKF3 | AT2G48010 | RKF3 | 1 | | | 2.8 | 0.4 | 0.4 | 0.9 | 0.9 | 1.2 |
| RKF3 | AT1G11050 | Y1105 | 2 | 625 | C357 | 2.2 | 1.7 | -1 | -2 | 0.7 | 1.5 |
| RLCK-IV | AT4G00330 | CRCK2 | 1 | | | 3.3 | 0.9 | 0.4 | 2 | 1.6 | 1.3 |
| RLCK-IXa | AT3G26700 | F4JDN8 | 1 | | | 2 | 1.4 | 2.1 | 2.7 | 1.8 | 0.9 |
| RLCK-VIIa-1 | AT1G20650 | PBL21 | 1 | | | 1.6 | 3.2 | 1.2 | 1.5 | 2 | 1.2 |
| RLCK-VIIa-1 | AT5G13160 | PBS1 | 1 | | | 2.7 | 2.2 | 3.6 | 4.8 | 3.2 | 3.2 |
| RLCK-VIIa-2 | AT5G02290 | PBL11 | 1 | | | 2.4 | 2.8 | 3.1 | 2.5 | 1.7 | 1.9 |
| RLCK-VIIa-2 | AT5G47070 | PBL19 | 1 | | | 2.1 | 2.8 | 1.2 | 3.4 | 1.5 | 1.2 |
| RLCK-VIIa-2 | AT3G09830 | PCRK1 | 1 | | | 1.6 | 1.4 | 1.4 | 2.6 | 1.6 | 2.2 |
| RLCK-VIIa-2 | AT5G03320 | PCRK2 | 1 | | | 2.6 | 1.4 | 1.9 | 1.2 | 2 | 2.2 |
| RLCK-VIIa-2 | AT2G17220 | PIX13 | 1 | | | 2.4 | 3.2 | 2.8 | 2.9 | 2.8 | 2.7 |
| RLCK-VIII | AT3G17410 | CARK1 | 1 | | | 2.9 | 3 | 3 | 3.3 | 3.2 | 2.8 |
| RLCK-VIII | AT1G48210 | F4HWU0 | 1 | | | 2.4 | 1.1 | 2.8 | 2.1 | 3 | 2.9 |
| RLCK-VIII | AT1G06700 | PTI11 | 1 | | | 1.7 | 1.4 | 1.8 | 1.9 | 1.7 | 1.9 |
| RLCK-VIII | AT2G30740 | PTI12 | 1, 2 | 366 | C6, C7 | 11 | 6.2 | 6.8 | 7.3 | 7.6 | 13 |
| RLCK-VIII | AT3G59350 | PTI13 | 1 | | | 1.4 | 1.4 | 1.7 | 1.8 | 1.7 | 1.6 |
| RLCK-VIII | AT2G47060 | Y2706 | 1 | | | 2.6 | 1.6 | 2.2 | 2.2 | 3.2 | 2 |
| RLCK-XII-1 | AT4G35230 | BSK1 | 1 | | | 1.7 | 1.8 | 1.8 | 1.5 | 1.7 | 2 |
| RLCK-XII-1 | AT5G46570 | BSK2 | 1 | | | 1.3 | 1.3 | 1.5 | 1.3 | 1.3 | 1.7 |
| RLCK-XII-1 | AT4G00710 | BSK3 | 2 | 489 | C192, C271 | 7.3 | 5 | 6.8 | 4.5 | 6.7 | 4.2 |
| RLCK-XII-1 | AT5G59010 | BSK5 | 2 | 489 | C50, C195, C343 | 6.1 | 3 | 2.6 | 4.9 | 4 | 2 |
| RLCK-XII-1 | AT1G63500 | BSK7 | 1 | | | 2.3 | 2.2 | 2.2 | 2.2 | 2 | 2.9 |
| RLCK-XII-1 | AT5G41260 | BSK8 | 1 | | | 2.5 | 2.7 | 3.4 | 2.9 | 3.3 | 3.1 |
| RLCK-XV | AT1G52540 | Q8LDB7 | 1, 2 | 350 | C8, C9, C103, C212, C292 | 8.4 | 3.3 | 6 | 3.8 | 6.2 | 6.5 |
| SD-2b | AT5G20050 | Y5005 | 1 | | | 2.5 | 0 | 1.4 | 0.7 | 0.8 | 1.4 |
| WAK | AT1G21270 | WAK2 | 1 | | | 1.7 | 1.1 | 1.6 | 1 | 1.2 | 1.3 |

**Figure S7.** Summary of S-acylated RLKs identified by both ABE-protein and ABE-peptide methods. Enriched folds in six genotypes are numerically indicated and highlighted with colors: red, enriched; blue, non-enriched.

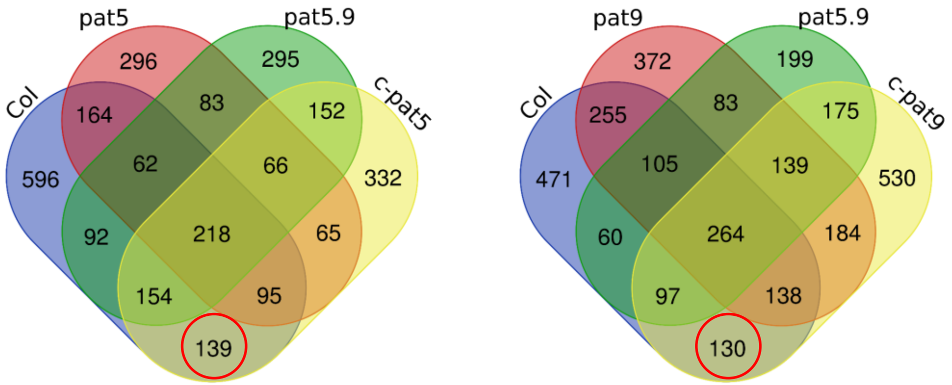


**Figure S8.** Venn diagram for S-acylated peptides identified in each genotype using the ABE-peptide method. Candidate substrates for PAT5 and PAT9 are indicated with red circles.

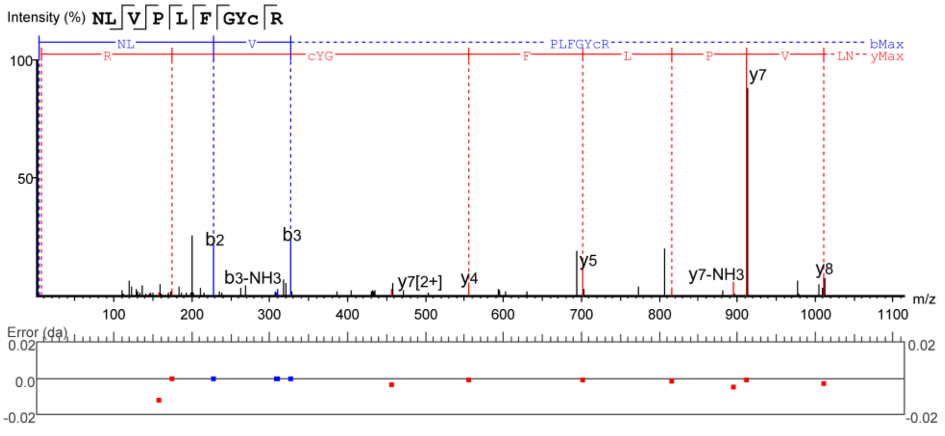


**Figure S9**. MS/MS spectrum of the S-acylated cysteine peptide from P2K1 identified by IP-MS based S-acylproteome: NLVPLFGYC(+57.02)R.
